# scDEcrypter: Uncertainty-aware differential expression analysis for viral infection in scRNA-seq

**DOI:** 10.64898/2026.03.09.710583

**Authors:** Luer Zhong, Karl Ensberg, Scott A. Tibbets, Aaron J. Molstad, Rhonda Bacher

## Abstract

Single-cell RNA-seq studies of viral infection are limited by sparse viral reads, under-labeled infected cells, and bystander responses that confound differential expression (DE) analysis. We introduce scDEcrypter, a penalized two-way mixture model that leverages partial infection and cell type labels to estimate latent cell-state probabilities. scDEcrypter provides fast permutation-based approximate p-values for prioritizing infection-associated genes while accounting for the uncertainty in cell-state. In simulations and applications to two viral infection datasets, scDEcrypter improved recovery of infected cell states and identified more biologically coherent infection-associated genes and enriched pathways.

## Background

Viral infections initiate complex and heterogeneous responses in the host as pathogens enter cells, replicate, and evade immune detection [1]. Understanding the variability of infection responses at the cellular level is critical for uncovering mechanisms of viral pathogenesis and host defense. Single-cell RNA sequencing (scRNA-seq) technologies have been successfully used to investigate cell type heterogeneity in infected tissues for a variety of viruses including influenza [2, 3], coronaviruses [4, 5], and flaviviruses [6, 7, 8]. However, one of the largest challenges in studying viral infections with scRNA-seq data lies in the ability to identify infected cells. viral scRNA-seq studies often have relatively few cells labeled as infected, ranging from < 1% to 5% in studies of single cell types infected in vitro [3, 8]. This percentage may increase over longer time courses and with higher multiplicities of infection; however, most scRNA-seq experiments suffer from under-identification of infected cells, largely due to difficulties in detecting and quantifying viral transcripts [9].

Although a few pipelines have emerged that improve specific virus quantification or enable entire virome profiling from sequencing data [10, 11], challenges remain. viral genomes mutate often or have sequences that are similar to those in the host genome [12], leading to viral reads being discarded or misaligned. Additionally, because of the sparsity and low sensitivity of scRNA-seq [13], viral transcripts may not be detected in cells with low levels of infection. Without detectable viral transcripts, reliably labeling infected cells is further complicated by so-called bystander cells, which exhibit similar transcriptional profiles as they respond to signals from infected cells [9]. As a result, downstream analyses have prioritized filtering out potential false positives, that is, cells that are not truly infected but contain a low number of viral transcripts associated with them due to contamination or through biological processes like phagocytosis [14]. These challenges in recovering infected cell labels have resulted in few cells being labeled as infected and have reduced the power to detect differential transcriptomic activity.

Without adequate apriori labels for each cell’s infection state, most differential expression (DE) methods for scRNA-seq are limited in their ability to distinguish infection-specific transcriptional changes [15, 16, 17]. Several approaches have been developed to accommodate uncertainty in cell labeling, including scANvI [18], miloDE [19], and GEDI [20]. However, these approaches all have limitations, particularly in addressing complex single-cell experiments that involve multiple variables (e.g., infection state and cell type). scANvI is able to annotate unknown cells along a single variable and performs DE testing only on genes utilized for the prediction. miloDE tests for DE genes within overlapping neighborhoods built from K-nearest neighbor graphs, but the outputs are often complex and their interpretation is less clear. GEDI models uncertainty of cell status based on known sample-specific factors; however, infected and uninfected cells must be present within the same sample.

To address the challenges in identifying infected cells and performing robust, interpretable inference across cell labels in scRNA-seq data, we developed scDEcrypter. Our method models infection status and other cell-level variables as latent variables with partial observability. We implement a regularized two-way mixture model, where mixture weights estimate cells’ probabilistic membership to combinations of cell states (e.g., infection status and cell type). The resulting weights are used to estimate cell-state-specific mean expression profiles and to account for cell-state uncertainty in differential expression analysis. We implement data-splitting to separate parameter estimation from approximate hypothesis testing. We demonstrate the accuracy of scDEcrypter through a comprehensive simulation study and compare its performance to existing approaches. In applications to an influenza and SARS-Cov-2 scRNA-seq datasets, scDEcrypter improved the recovery of infection states, identified infection-specific genes, and revealed trends in infection status over time.

## Results

### Overview of scDEcrypter

The workflow of scDEcrypter starts with pre-processing, followed by model training, estimation of cell-state weights, and differential expression testing (Figure 1). scDEcrypter requires the raw counts from scRNA-seq data, partial infection status labels, and an additional partitioning variable label, typically representing cell type. The additional partitioning variable may be fully or partially known and is generalizable to any cell-specific label. Once the data is split into a generation (training) and test set, the resulting two datasets are independently subject to normalization and a variance stabilizing transformation. For model training and estimation, the generation dataset is further reduced to a subset of highly variable features. A two-way mixture model over the latent cell states is fit using a variation of the expectation-maximization algorithm. After estimation, the trained parameters are used to infer cell-state weights in the test dataset for infection status and cell type. Finally, scDEcrypter performs cell type-specific differential expression testing between infection states, with the weights accounting for each cell’s state uncertainty. Detailed descriptions of the method are in the Methods section and in Additional file 1: Supplementary Methods.

**Figure 1.**
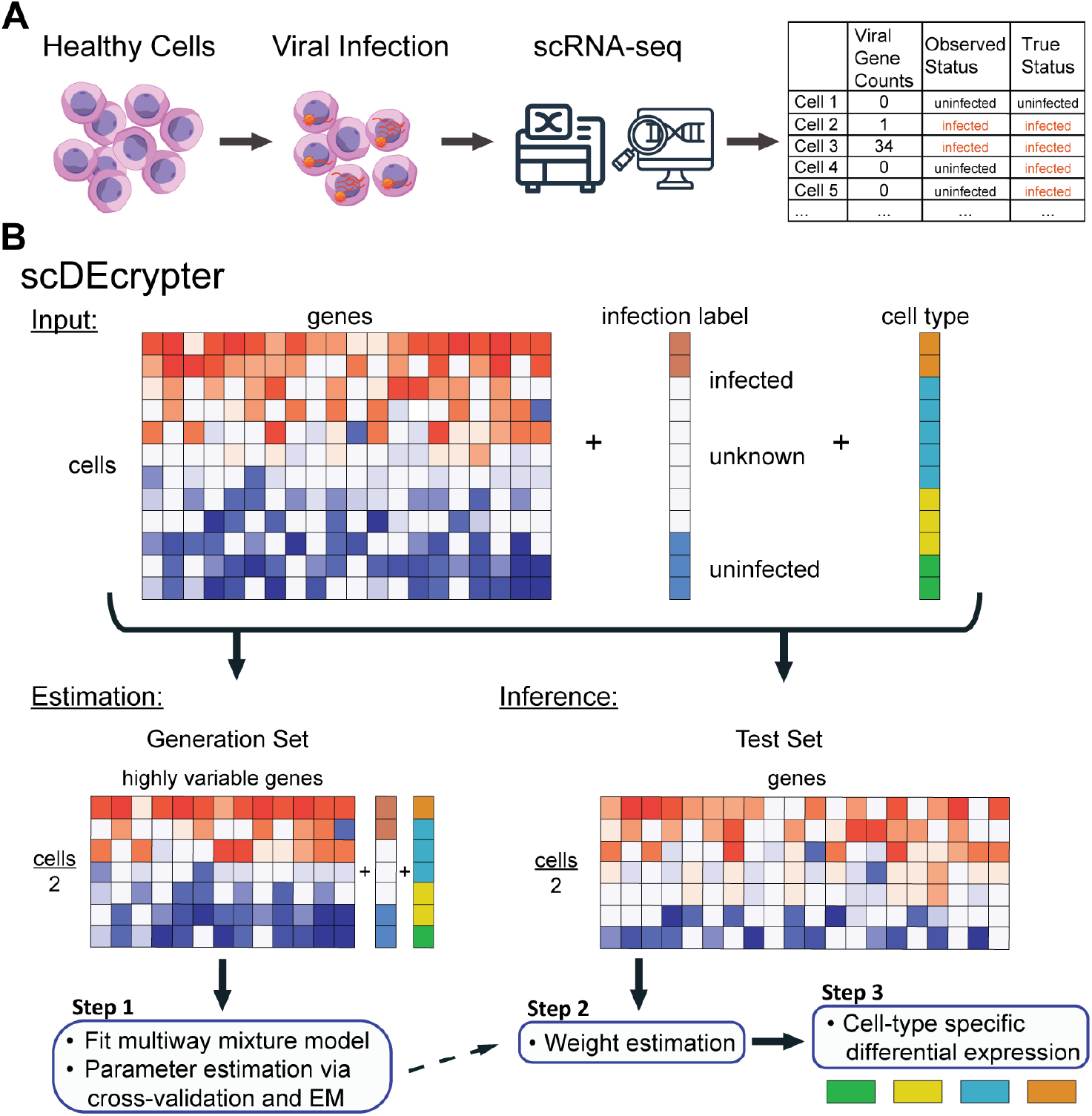
Overview of the scDEcrypter method. **A.** In viral scRNA-seq datasets, infection status is partially known. **B**. The input to scDEcrypter includes scRNA-seq data, partial infection labels, and labels for an additional cell-specific partitioning variable (e.g., cell type). After data splitting, the generation set is reduced to a subset of highly variable genes. Cross-validation is used to select tuning parameters, followed by model estimation. On the cells in the test set, the estimated parameters are used to infer cell weights, followed by a permutation test for differential expression testing.

### Accurate inference with scDEcrypter across partial-labeling simulations

We simulated scRNA-seq data using an extended version of the scaffold R package [21] to reflect the heterogeneity of viral infection at the cell type and sample level (details in Methods). The simulations were varied in the proportion of cells pre-labeled, the proportion of cell type associated genes, the proportion of infection associated genes, and the magnitude of the association for infection associated genes.

In practice, the weights inferred by scDEcrypter for the test set are used directly for differential expression testing, however, they may also be thresholded to assign cells to discrete infection states. For the simulations, we first evaluated how accurately scDEcrypter could predict infection state on unlabeled cells across varying proportions of cells pre-labeled for infection state, assuming cell type was fully known. Across all simulations, scDEcrypter achieved an overall balanced accuracy of 87.9%. In the moderate and large fold-change scenarios, this increased to 94.0% (Figure 2A). For subtle fold-changes, accuracies were expectantly lower, averaging 75.7%, but increased as the proportion of pre-labeled cells increased. Prediction accuracy was also influenced by the proportion of simulated infection-associated genes (Figure 2B). Overall, the poorest performing scenarios were with combinations of subtle fold-changes, pre-labeling limited to 1% of cells, and only 1% of genes associated with infection. This was most evident in the positive infection calls, where the positive predictive value was lowest under those scenarios (Additional file 1: Fig. S1).

**Figure 2.**
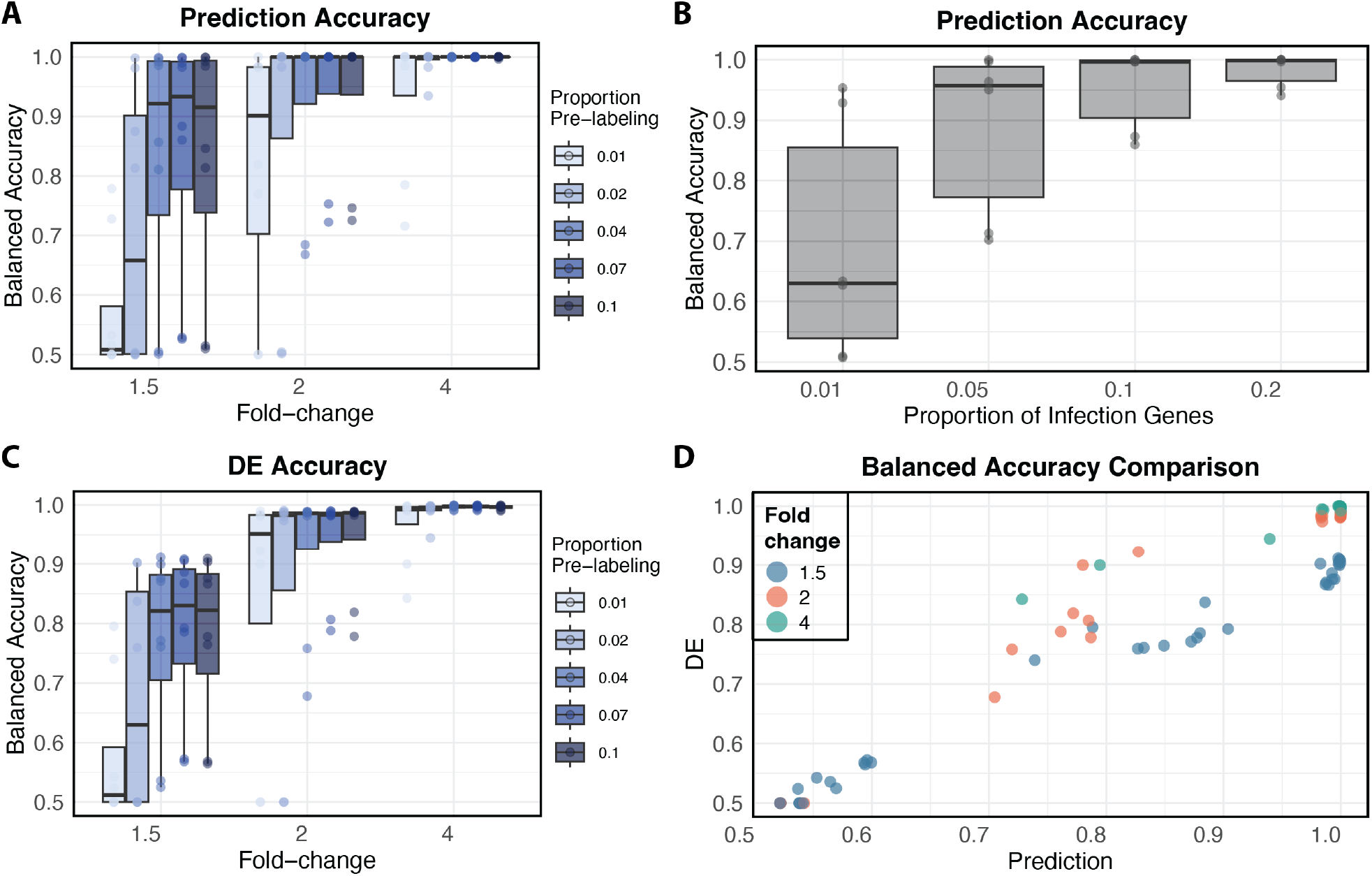
Simulation study results. **A.**Balanced accuracy for infection state prediction by the foldchange of infection-associated genes. Boxplots are colored by the proportion of pre-labeled cells for viral infection. **B**. Balanced accuracy for infection state prediction by the proportion of infection-associated genes. **C**. Similar in structure to (A) for the balanced accuracy for differential expression (DE) testing to identify infection-associated genes. **D**. The relationship between the balanced accuracy for differential expression testing and prediction across all simulation scenarios.

In differential expression testing, where the inferred cell weights are used directly, scDE-crypter achieved an average balanced accuracy of 94.5% in identifying infection-associated genes at moderate and high fold-changes (Figure 2C). At a more subtle fold-change of 1.5, balanced accuracy for DE testing increased with a higher proportion of pre-labeled cells (Figure 2C). Inferential accuracy was highly correlated with prediction accuracy, with higher prediction accuracies leading to higher DE accuracy (Figure 2D).

To highlight one of scDEcrypter’s key innovations in accounting for uncertainty in cell-state based on a small subset of confidently labeled cells, we compared its performance to widely used DE testing methods for scRNA-seq data. Cell’s infection state labels for the comparison methods (Seurat [15], MAST [16], and DESeq2 [17]) were set to mirror the settings used in standard analysis practice, with confidently labeled cells (those pre-labeled in scDEcrypter) grouped as infected and all other cells set to uninfected. Across pre-labeling and gene infection proportion settings, scDEcrypter consistently achieved the highest balanced accuracy (Figure 3A) and F1 scores (Figure 3B) in identifying infection-associated genes. DESeq2 performed only slightly worse on average, but showed a much greater variability across simulations compared to scDEcrypter. Seurat and MAST demonstrated less reliable performance with reduced robustness across simulation scenarios.

**Figure 3.**
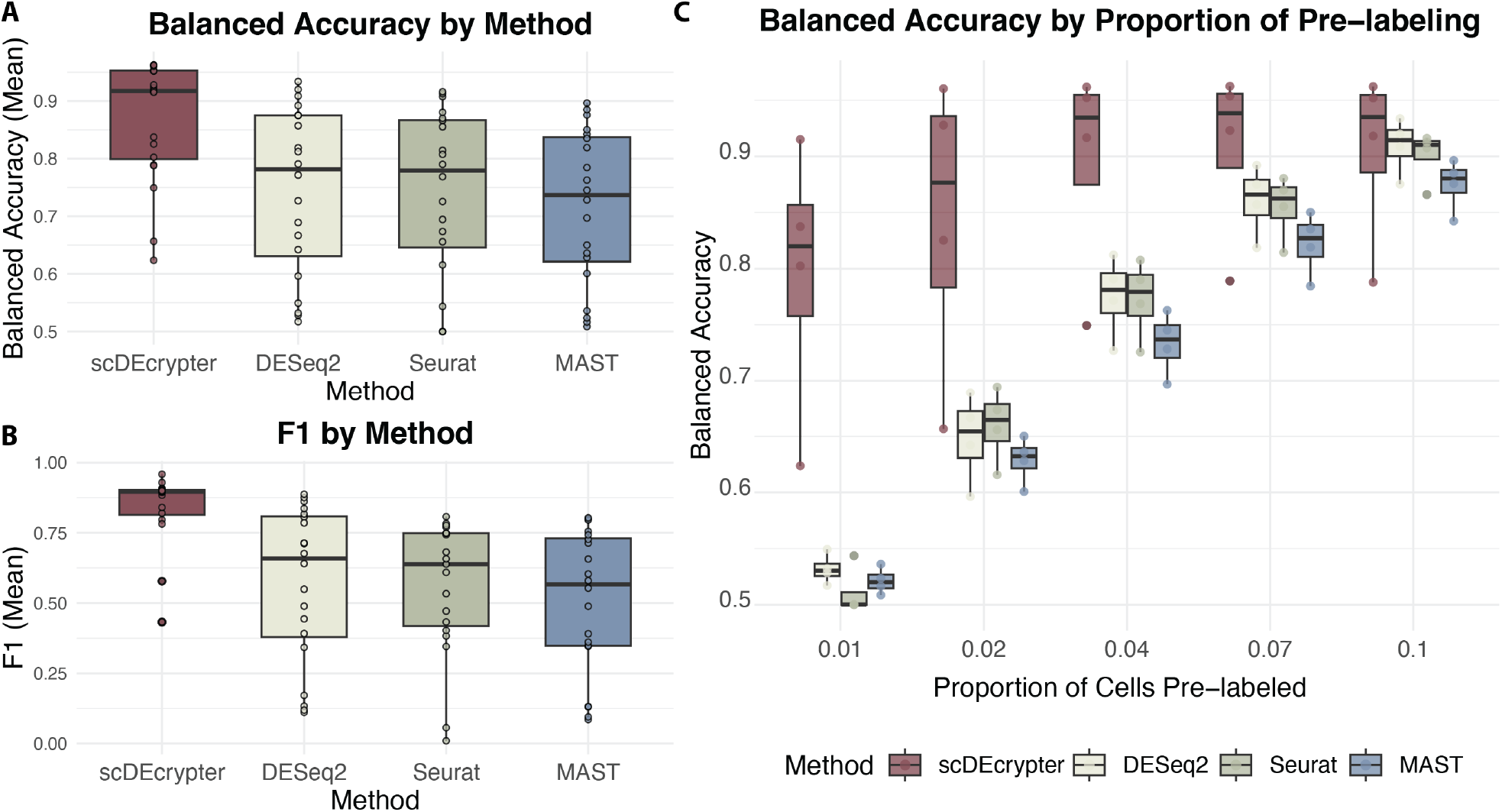
Comparison to other DE testing methods. **A.**Balanced accuracy averaged over simulated scenarios for four DE testing methods: scDEcrypter, DESeq2, Seurat, and MAST. Accuracy was averaged across all replicates, fold-changes, the number of HvGs, and cell type proportions. **B**. F1 scores across methods. **C**. Balanced accuracy across varying proportions of pre-labeled cells (x-axis). Methods are distinguished by color.

We also compared scDEcrypter to scANvI [18], a semi-supervised deep generative model (Additional file 1: Fig. S2). scANVI was evaluated under the same simulation scenarios and train/test splits (see Methods). Because scANvI requires labeled cells for each infection state, we pre-labeled comparable subsets of infected and uninfected cells within each cell type; all remaining cells were treated as unlabeled during model fitting. Model fitting was restricted to training cells, whereas infection-state prediction and DE detection were evaluated in held-out test cells. scDEcrypter achieved higher balanced accuracy and F1 scores for infection-state prediction across signal settings. For DE detection, scDEcrypter showed consistently higher balanced accuracy, while F1 scores were largely similar between the two methods.

We next evaluated the performance of scDEcrypter when both cell state variables were not fully observed, that is having both infection state and cell type as partially labeled. In addition to the labeling schemes we used for viral infection for the simulation, we also randomly labeled only 75% of cells in each cell type. The overall balanced accuracy for infection state prediction on unlabeled cells averaged 82.1%, comparable to the results obtained when cell types were fully observed (Figure 4A). The balanced accuracy for cell type prediction was consistently above 90% across settings (Figure 4B).

**Figure 4.**
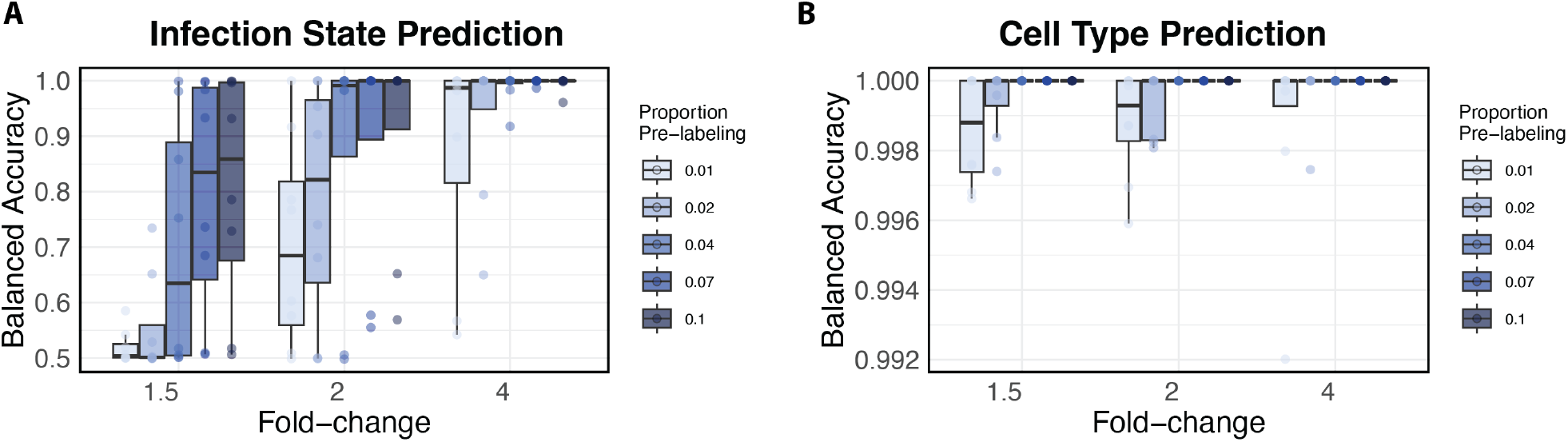
Simulation results with partial observation of both cell state variables. **A.**Balanced accuracy for infection state prediction by the fold-change of infected associated genes; boxplots colored by the proportion of pre-labeled cells for viral infection. **B**. Balanced accuracy for cell type prediction by fold-change of infected-associated genes. Boxplots are colored by the proportion of pre-labeled cells for viral infection.

We also assessed scDEcrypter’s computational performance by benchmarking runtime and peak memory usage while varying the number of cells, genes, and cell types (Additional file 1: Fig. S3). Runtime and memory were largely insensitive to the number of cell types and scaled favorable with dataset size. For a dataset of 20,000 cells, model estimation using 200 training genes completed in approximately 3.6 minutes, and permutation-based DE testing of 5,000 genes completed in 2.7 minutes using eight CPU cores. Median peak memory usage remained below 3 GB for this dataset size. Across benchmark settings containing up to 20,000 cells, peak memory usage was typically 1 to 3 GB, increasing to approximately 10 GB for the largest datasets evaluated (100,000 cells).

Additional simulations examining the robustness of scDEcrypter to model misspecification and violations of key assumptions are described in Additional File 1: Simulation studies under model misspecification.

### scDEcrypter identifies influenza infection-associated genes

Next, we evaluated scDEcrypter’s performance on a dataset where infection state was the only cell state of interest. An influenza time-course experiment was carried out by Russell et al. using a homogeneous population of cells derived from a single cell line [3]. Cells were collected for single-cell RNA sequencing at four time points: 0(uninfected)-, 6-, 8-, and 10-hours post infection [3]. To mitigate potential batch effects, sample time was included as a fully known partitioning variable with unique labels assigned to the two replicates of the 8-hour time point. Across all time points, influenza mRNA was detected in approximately five percent of cells. In the original study, 368 cells were labeled as confidently infected based on an empirically defined threshold [3]. To reflect practical application of our method, we ran scDEcrypter with cells pre-labeled as infected according to the original publication (n=368), and cells in the 0-hour sample pre-labeled as uninfected.

Applying a strict threshold to the weight matrix inferred by scDEcrypter, we found 17.4% of the unlabeled cells in the test set as highly likely to be infected. This sharply contrasts with the approximately 5% of infected cells identifiable based solely on viral mRNA counts[3]. However, our estimate aligns more closely with the multiplicity of infection (MOI) of 0.3 used in the study, which the authors reported as an estimated infection rate upper-bound of 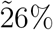 [3]. We also observed that the proportion of cells predicted as infected varied depending on the probability threshold used (Additional file 1: Fig. S4), noting that scDEcrypter’s advantage is that it avoids arbitrary cutoffs and instead uses the weights directly for downstream inference.

Next, scDEcrypter was used to test for differential expression between infected and un-infected cells at each time-point (Supplemental Table 1). We compared the results of the differential expression testing to those obtained via Seurat [15]. Overall, scDEcrypter identified 2,195 shared DE genes across all time points. In comparison, Seurat identified only five human genes that were shared between any two time-points and only the eight influenza viral transcripts were DE across all samples (Figure 5A&B). We also compared our results to those using scANvI [18], which leverages partial cell type annotations to infer labels for unannotated cells. Out of 5,000 genes evaluated for DE, scANvI was only able to predict 360 DE genes total, of which only 13 were DE at all time points. Because our simulations indicate that scDEcrypter p-values can be anticonservative when infection intensity varies across cells (see Discussion and Additional File 1), the scDEcrypter DE set likely includes some false positives. We thus focus below on genes and pathways supported consistently across time points and by external gene sets.

**Table 1.** Overview of all simulation scenarios.

| Prop. Celltype genes | Prop. Infection genes | Infection fold-change | Prop. Pre-labeling |
| --- | --- | --- | --- |
| .05 | .01 | 1.5 | .01 |
| .2 | .05 | 2 | .02 |
|  | .1 | 4 | .04 |
|  | .2 |  | .07 |
|  |  |  | .1 |

**Figure 5.**
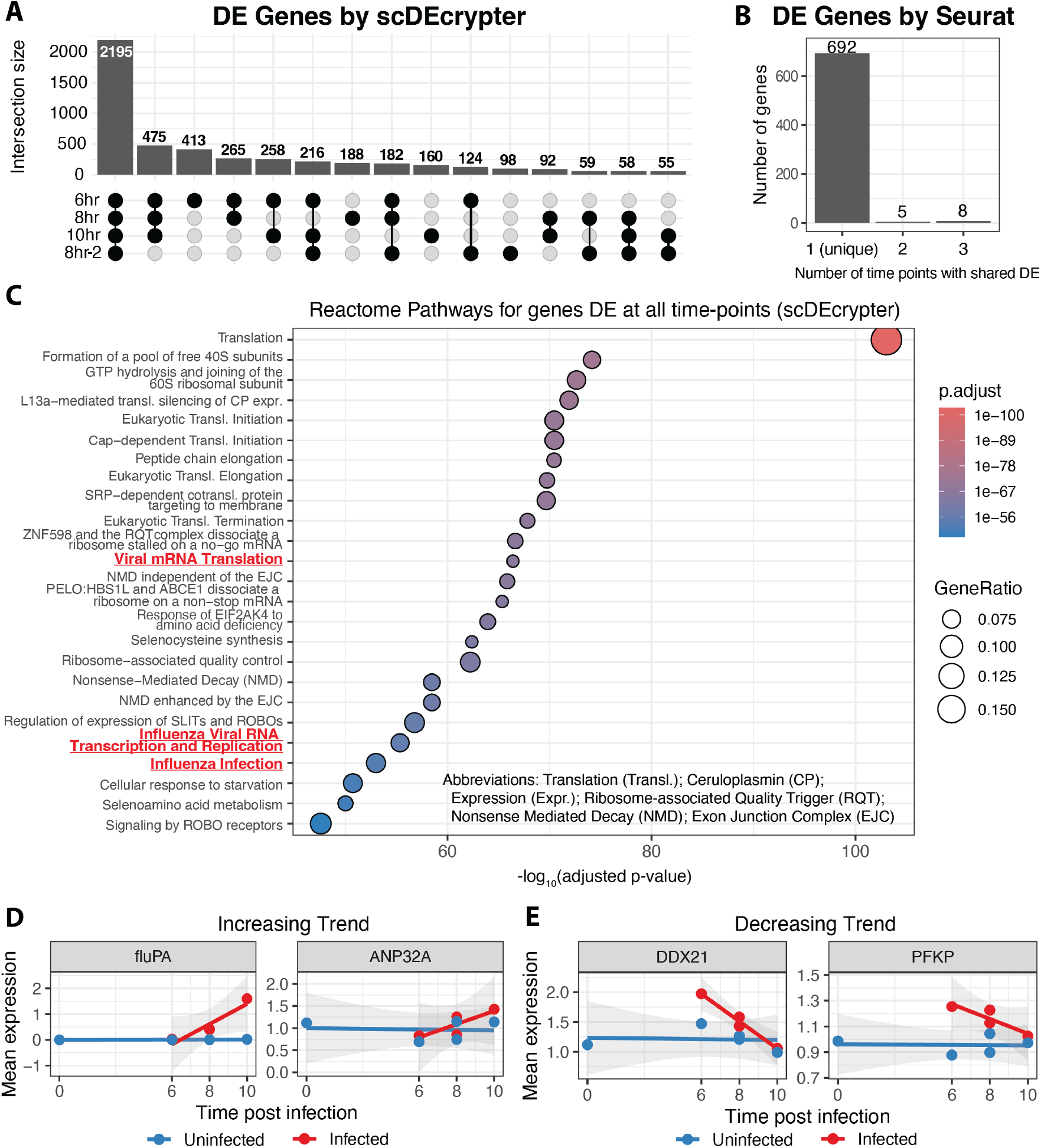
Results on the influenza dataset. **A.**Upset plot of DE genes identified by scDEcrypter. **B**. DE genes identified by Seurat, grouped by the number of time points at which they are DE. Uniquely DE genes are present in only one time point. Only influenza genes were DE in all time-points. **C**. Pathway enrichment of genes consistently identified by scDEcrypter as DE across all time-points. **D**. Two representative DE genes from scDEcrypter with an increasing mean expression trend in the infected state. The sample time is shown on the x-axis and the y-axis is the gene mean estimated by scDEcrypter. **E**. Similar to (**D**) for representative genes with a decreasing mean expression trend in the infected state.

To further validate our findings, we compared our results to the original list of DE genes by Russel et al. [3], our method identified all but three of the 42 genes tested as differentially expressed, while Seurat only identified approximately half (56%). We also examined our ability to identify genes associated with direct viral infection using a gene list curated by Watanabe et al. [22]. Of the 60 highly curated viral replication-associated genes tested, scDEcrypter identified 55 (92%) as DE and Seurat identified only 13 (22%).

Pathway enrichment analysis of the shared DE genes identified by scDEcrypter also clearly captured the biology of viral replication. Enrichment was dominated by translation- and ribosome-related pathways, as well as influenza infection pathways, directly reflecting influenza’s reliance on hijacking host protein synthesis (Figure 5C).

We next investigated temporal expression trends by identifying genes with progressive changes (increasing or decreasing) over the infection time course based on state-specific gene means estimated by scDEcrypter (Supplemental Table 2). All influenza genes showed increased expression in the infected state over time and their expression levels at the early (6-hr) time-point was similar to that in the uninfected state, reflecting the onset and sub-sequent progression of viral replication (representative gene shown in Figure 5D). Human genes with progressively increasing trends were generally enriched for protein catabolism, ubiquitin-proteasome degradation, and antigen processing (Additional file 1: Fig. S5A). Among the increasing genes was *ANP32A* (Figure 5D), which encodes a host factor required for influenza A virus polymerase activity and viral RNA replication [23, 24]. Genes with progressively decreasing expression trends in infected cells also showed enrichment for RNA processing/splicing and negative regulation of programmed cell death (Additional file 1: Fig. S5B). One of these genes, *DDX21* (Figure 5E), encodes an RNA helicase that restricts influenza A virus replication by binding PB1 and inhibiting viral polymerase activity; its decrease may weaken antiviral restriction in infected cells [25]. Separately, *PFKP*, one of the most strongly decreased genes, encodes a key glycolytic enzyme, and influenza-associated glycolytic reprogramming has been linked to viral replication and inflammatory responses [26].

### scDEcrypter distinguishes infection and bystander transcriptional signals in SARS-Cov-2

We next applied scDEcrypter to a more heterogeneous scRNA-seq dataset consisting of multiple human bronchial epithelial cell types cultured in vitro and experimentally infected over a three-day time course [14]. Approximately 72% of all virus-exposed cells had at least one SARS-Cov-2 viral read, corresponding to 21%, 84%, and 88% of cells at one, two, and three days post-infection, respectively. To account for ambient virus or read misalignment, the original analysis used a threshold of ten viral reads, which resulted in an overall infection rate of 8.6% (over 4,500 cells). Given the large number of cells available for pre-labeling, we aimed to use scDEcrypter to characterize biological heterogeneity among virus-exposed cells by considering three possible infection states: infected cells, bystander cells exhibiting reactive responses, and cells resembling the uninfected control cells.

For pre-labeling, cells from the uninfected sample were pre-labeled as uninfected, and any cells with more than 40 SARS-Cov-2 viral reads were pre-labeled as confidently infected. The conservative pre-labeling threshold was chosen as a handful (n=27, < 1%) of cells in the uninfected sample had viral reads present, with one control cell having as many as 31 SARS-Cov-2 viral reads. Cell type was included as the additional partitioning variable, using annotations from the original analysis. Cells were defined among eight epithelial cell types as: basal cells, BC/club, ciliated cells, club cells, goblet cells, ionocytes, neuroendocrine cells, and tuft cells. We ran scDEcrypter with cell type as partially latent on the generation set (75% pre-labeled) to allow for additional information sharing across cell types regarding infection state at the estimation stage.

As this dataset included an additional partitioning variable (cell type), we first explored how cell state prediction was affected by varying levels of pre-labeling. We applied a threshold to the weights inferred by scDEcrypter to predict cell state. We observed a linear decrease in cell type prediction accuracy by about 2% for every 5% reduction in pre-labeling in the test dataset (Additional file 1: Fig. S6). Since partial labeling in the inferential stage offered no advantage when all cell type labels were available in the test dataset, we proceeded by using all cell type labels in our analyses. As a further validation of test-set weight estimation, we cleared pre-labeling of infected cells on the test dataset and estimated each cell’s probability of belonging to the uninfected state. This probability decreased sharply with increasing SARS-Cov-2 mRNA count (Additional file 1: Fig. S7). We then applied a threshold to the weights to examine infection state, finding that scDE-crypter predicted an increasing proportion of infected cells post infection: 14% at one day, 17% at two days, and 25% at three days (Figure 6A). Although the authors used MOI 0.01 for the inoculation, the infection rates observed here are consistent with the 10-12 hour replication rate of SARS-Cov-2 and the ability of each infected cell to generate 10^3^–10^4^ new infectious virus particles per cell.

**Figure 6.**
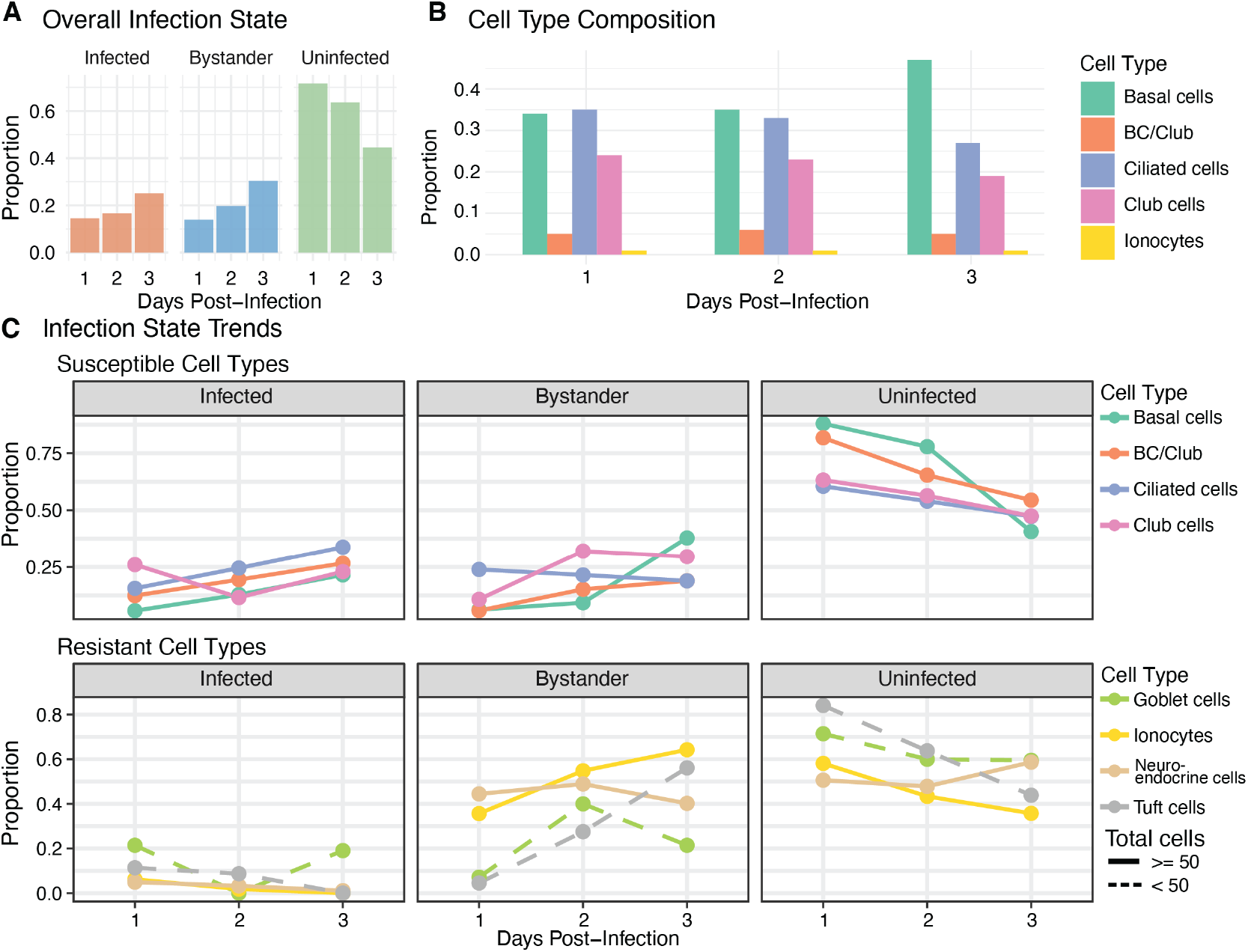
Predictive assessment of SARS-Cov-2 infection states. **A.**Proportion of all cells belonging to each infection state over days post-infection after thresholding the scDEcrypter weight matrix. **B**. Cell type composition over days post-infection. Goblet, neuroendocrine, and tuft cells have proportions rounding to zero and are not shown. **C**. Cell type specific infection state trends over days post-infection. Cell types were split into susceptible (top) or resistant (bottom) groups based on their infection trends and the original publication. Dashed lines indicate cell types having fewer than 50 total cells present in the dataset.

We next examined how the predicted infection and bystander states were distributed among specific cell types and time points. By three days post-infection, the cells that were engaged with the virus as either infected or bystander comprised the majority of the population (Figure 6A). Day three also marked a shift in cell type composition, with an increase in basal cells and decreases in both ciliated and club cells (Figure 6B). Even as the cell type composition remained stable over days one and two, the ciliated, BC/club, and basal cells showed increasing proportions of infected cells over time (Figure 6C). Ciliated cells had the largest number of cells at one day post-infection engaged with SARS-Cov-2 as either infected or bystander. Club cells were the most infected cell type at one day post-infection, but appeared to shift predominantly to the bystander state by day two. Consistent with the original study, goblet, ionocyte, neuroendocrine, and tuft cells appeared more resistant to infection. Few cells were predicted to be infected and their proportions remained relatively stable across the time course (Figure 6C), though they appear to be increasingly engaged as bystander cells in days two and three. However, these cell types collectively represent less than 2% of all cells in the dataset making it difficult to confidently quantify their infection dynamics.

We next used scDEcrypter to identify transcriptional differences between the three infection states by performing cell type-specific DE analysis on infected versus bystander and bystander versus uninfected states for each time point (Supplemental Table 3). Ciliated cells had the most DE genes at all time points when comparing infected versus bystander cells (Figure 7A). In the bystander versus uninfected comparison, BC/club cells had the largest number of DE genes at one day post-infection, basal cells had the most at day two, and ciliated cells had the highest at day three. For genes that were DE in at least four of the eight cell types at any day, we performed enrichment analysis on genes split into three groups: genes exclusively DE in the infected versus bystander comparison, genes exclusive to the bystander versus uninfected comparison, and genes that were DE in both comparisons (Figure 7B). Genes DE in both comparisons were largely enriched for translation-related activities. Genes specific to the bystander versus uninfected comparison were enriched for pathways related to cellular energy production (mitochondrial ATP synthesis, oxidative phosphorylation, electron transport chain), activation of stress responses, and immune surveillance. Genes uniquely DE in the infected versus bystander comparison were enriched for viral infection pathways, including SARS-Cov infection specifically. Among genes contributing to SARS-Cov-associated Reactome pathways, eight (*RPS27L, HN-RNPA1, YWHAE, PCBP2, SFN, ITGB1, ATP1B3*, and *CSNK1A1*) were consistently differentially expressed only between infected and bystander cells across at least four cell types. Several of these genes have established linked to SARS-Cov-2 biology, including *HNRNPA1*, which has been implicated in viral RNA regulation [27], *ITGB1*, which participates in spike-associated host responses [28], and *YWHAE* and *SFN*, members of the 14-3-3 protein family that interact with the SARS-Cov-2 nucleocapsid protein [29].

**Figure 7.**
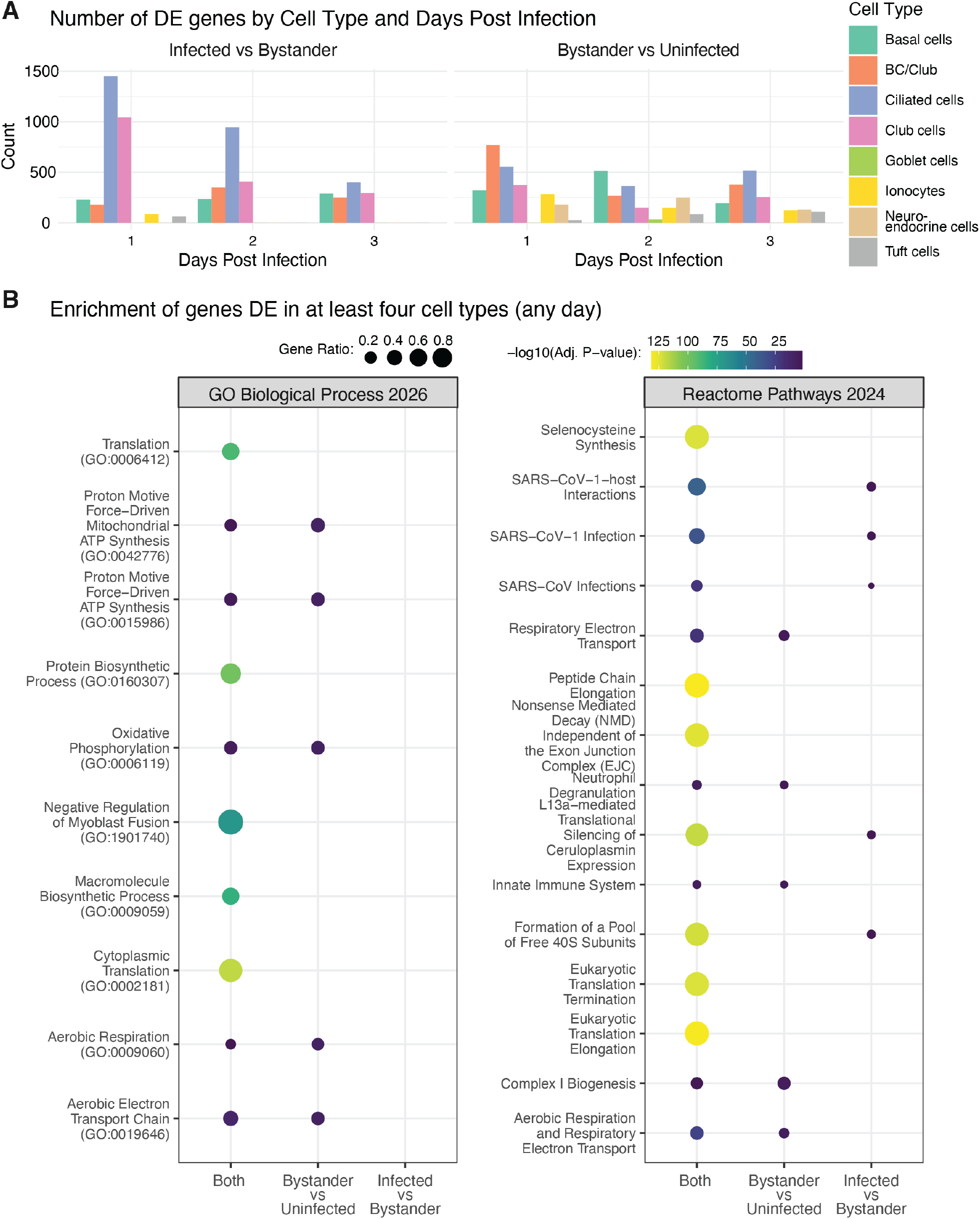
Differential expression between SARS-Cov-2 infection states. **A.** The number of DE genes identified by scDEcrypter for each cell type and time-point for infected versus bystander (left) and by-stander versus uninfected (right) comparisons **B**. GO Biological Processes and Reactome Pathways enrichment of genes DE in at least four cell types at any time-point exclusive to bystander versus infected comparison, exclusive to infected versus bystander comparison, or DE in both comparisons.

## Discussion

We introduce scDEcrypter, a statistically principled framework for improved differential expression analysis in scRNA-seq data with partially latent states. scDEcrypter implements a penalized multiway mixture model that leverages partial labels to anchor latent states and employs a data-splitting strategy to improve downstream testing. By combining permutation-based approximate inference with a small set of confidently labeled cells, scDEcrypter infers cell-state weights and tests for infection-associated transcriptional changes.

Under-labeling is a common limitation in viral scRNA-seq studies, especially in clinical samples where the proportion of infected cells is likely to be even lower due to greater cellular heterogeneity and variation in sample collection. Unlike most current single-cell DE methods that require fully specified groups, scDEcrypter relaxes this assumption by allowing infection status and an additional partitioning variable to be partially latent. Our extensive simulations and applications to influenza and coronavirus datasets demonstrated that scDEcrypter improved the power to detect infection-associated genes and revealed biologically coherent pathways. The enriched pathways identified by scDEcrypter align closely with known viral biology, underscoring the biological interpretability of our method’s outputs.

Despite these strengths, there are some limitations. The current model assumes Gaussian-distributed stabilized expression values, which may not fully capture the overdispersion characteristic of scRNA-seq data. While the variance stabilizing transformation helps to address this, more flexible distributional assumptions may be beneficial. The permutation-based p-values do not require Gaussian expression, but their validity depends on the exchangeability conditions specified in the Methods; equality of means alone is insufficient. In Additional File 1: Simulation studies under model misspecification, we show that our test can maintain the nominal level even when the data are generated from a negative binomial rather than Gaussian, provided infection effects are homogeneous across infected cells (Table R2). The simulation studies in Additional File 1 also indicate that when infection intensity varies across infected cells, particularly when cells also share strong latent factors inducing correlation, our permutation test’s p-values can be anticonservative, so p-values are best used to rank genes and FDR-adjusted calls should be interpreted as liberal. Additionally, the data-splitting strategy, though effective in reducing bias, may limit power in small datasets with few labeled infected cells. Finally, scDEcrypter currently models infection status and cell type independently, whereas infection can reshape cellular identity; future extensions could incorporate these dependencies to further enhance DE accuracy.

A related assumption of our model is that, conditional on cell type (or the specified partitioning variable), pre-labeled cells are representative of the latent state to which they are assigned. Thus, labeled cells serve as anchors for inference in unlabeled cells, and inferred infection probabilities may reflect similarity to these anchors rather than the probability of infection in an absolute sense. This consideration is particularly relevant when labels are defined using high-confidence viral mRNA evidence or experimental time points, since such cells may not represent the full biological spectrum of infection. The model can accommodate one additional latent state beyond those represented by the pre-labeling; however, distinct latent states or systematic expression shifts not represented by any labeled cells cannot generally be detected reliably or assigned a biologically interpretable meaning. We therefore recommend that users evaluate, using domain knowledge, whether their labeling scheme provides anchors for all biologically plausible states before applying scDEcrypter. Even when this assumption is imperfect, the model can provide a useful alternative to applying a uniform viral-read threshold, which does not account for biological heterogeneity across cells.

## Conclusion

scDEcrypter offers a robust and interpretable approach for identifying cell type-specific factors critical to viral tropism, replication, and immune evasion. Its flexible framework may also be applicable to other biological contexts with partially latent group labels, as long as some confidently labeled cells are present, such as cancer subclones, drug-resistant cell populations, or immune activation states. Explicitly modeling uncertainty in cell-state membership provides a powerful solution to the limitations imposed by sparse viral reads and under-labeling in single-cell studies of viral infection.

## Methods

### Preliminaries

The scDEcrypter approach assumes that given the *i*-th cell’s cell type (*C*_*i*_) and viral status (v_*i*_), the *j*-th gene’s expression satisfies

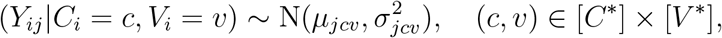

where here, 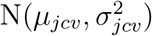 denotes a normal distribution with mean *µ*_*jcv*_ ∈ ℝ and variance 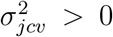. Throughout, we use [*C*\*] := {1, 2, · · ·, *C*\*} and [*v*\*] := {1, 2, …, *v*\*} to denote the discrete set of possible cell types and viral statuses. The additional partitioning variable *C*_*i*_ does not have to correspond to cell type, but could be another cell-specific label. This approach assumes that marginally, *Y*_*ij*_ follows a mixture of normals over all (*c, v*) ∈ [*C*\*] × [*v*\*], with mixing parameter *π*_*cv*_ = Pr(*C*_*i*_ = *c, V*_*i*_ = *v*). scDEcrypter estimates 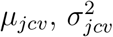, and *π*_*cv*_ using penalized maximum likelihood, employing a variation of the expectation-maximization algorithm. Details are described below and in Additional file 1: Supplementary Methods. The scDEcrypter method is available as an R package on GitHub at https://github.com/bachergroup/scDEcrypter. We describe the inputs and running of scDEcrypter below. A table summarizing key parameters and providing best practice guidance in Supplemental Table 4.

### Pre-processing

Before fitting the model, the scDEcrypter framework requires several pre-processing steps of the scRNA-seq data. First, scDEcrypter requires a subset of cells to be pre-labeled based on observed viral signals to help anchor latent infection states during model inference. We recommend assigning infection status using empirical criteria that reflect strong or absent infection signals, such as reads aligned to the viral genome or cells from an un-infected control. Next, the data is partitioned into two sets: the generation set and the test set. This split ensures that the same data are not used for both estimating mixture parameters and testing hypotheses. The remaining pre-processing including normalization and transformation will be performed separately on the two datasets to prevent data leakage. The partitioning is done with respect to the *C*_*i*_ and v_*i*_ labels (when possible) to ensure representative model training. On each set, normalization and variance stabilizing transformation pre-processing steps are done independently. The counts are normalized and variance-stabilized using the `transformGamPoi` R package v1.10.0[30], applying the shifted logarithm transformation with a shift of one and using the “poscounts” option to estimate the scale factors. The shifted-log transformation closely approximates the delta method variance-stabilizing transformation for Gamma–Poisson distributed data and effectively addresses heteroskedasticity [30].

For training the model, we select a fixed number of highly variable genes (HvG) to reduce dimensionality and focus the model on the most informative features. Genes are selected based on their gene-wise variance calculated after variance stabilization on the generation set. This is done with respect to the *C*_*i*_ when appropriate, and the HvG set is taken to be the most variable genes as the compact training panel. From the same ranking, a larger panel of genes is also defined for downstream analysis. Importantly, we apply different HvG numbers for the estimation and inferential stages to balance model efficiency and downstream power. Specifically, the generation set uses a small number of HvGs (e.g., 100 genes) to reduce overfitting and encourage stable parameter estimation during model training. In contrast, the inferential stage employs a substantially larger number of HvGs (e.g., 5,000 genes) to maximize resolution for differential expression analysis and improve detection of subtle transcriptional differences between inferred infection states. This allows scDEcrypter to remain computationally tractable during model fitting, while fully leveraging the richer gene set during downstream analysis.

Through this carefully structured pre-processing pipeline, scDEcrypter is able to reduce technical noise, enhance biological signal, and initialize latent structure learning with minimal (subjective) supervision, thereby enabling more accurate modeling of infection states and downstream differential expression analysis.

### Multiway mixture model and penalized maximum likelihood estimator

Assuming the generation dataset contains *n* cells and *d* genes, the probability density function for *Y*_*ij*_ given *C*_*i*_ = *c, V*_*i*_ = *v* is

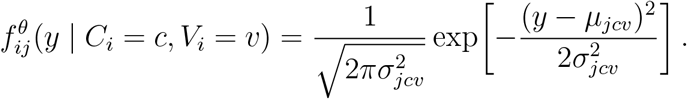

We abbreviate this conditional density as 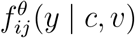. Let *θ* collect all model parameters:

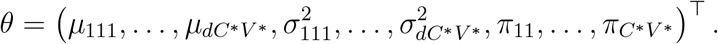

Thus, marginally, *Y*_*ij*_ follows a mixture of normals, i.e., has density

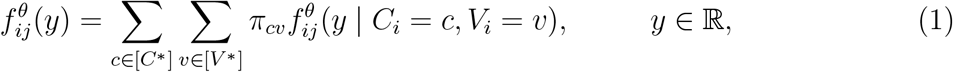

where *π*_*cv*_ = Pr(*C*_*i*_ = *c, V*_*i*_ = *v*) for each (*c, v*) ∈ [*C*\*] *×* [*v*\*].

As mentioned, we typically do not observe realizations of both *C*_*i*_ and *v*_*i*_ for each *i* ∈ [*n*]. Thus, when constructing the observed data log-likelihood, we must account for which variables are observable for the *i*-th subject. When neither *C*_*i*_ nor *v*_*i*_ is observed, the observed data likelihood contribution is the likelihood of *Y*_*i*_. When only *C*_*i*_ is observed, the likelihood contribution is the joint likelihood of (*Y*_*i*_, *C*_*i*_), and similarly when only *v*_*i*_ is observed. When both *C*_*i*_ and *v*_*i*_ are observed, the likelihood contribution is the joint likelihood of (*Y*_*i*_, *C*_*i*_, *v*_*i*_). We assume that *Y*_*ij*_ is independent of *Y*_*ij*_′ for all *j ≠* (*C*_*i*_, *v*_*i*_). Thus, to write the observed data log-likelihood, define

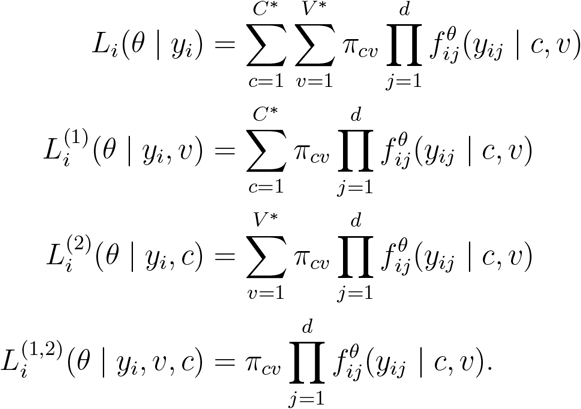

Let *B* be the index set of cells for which we observed both *C*_*i*_ and *v*_*i*_, let *C* (resp. *v*) be the set for which we observe only *C*_*i*_ (resp. *v*_*i*_), and let *N* be the set for which we observe neither *C*_*i*_ nor *v*_*i*_. Hence, the observed data log-likelihood can be expressed

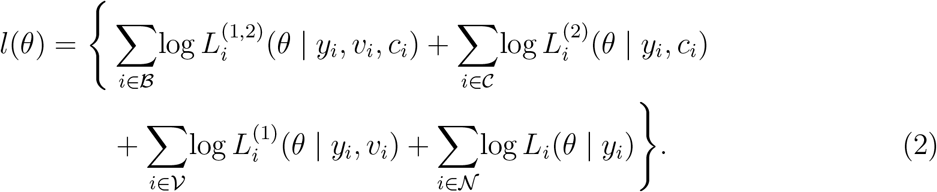

The unpenalized maximum likelihood estimator is thus arg max_*θ*_ *l*(*θ*). Because *d* is still relatively large, we propose a penalized maximum likelihood estimator that regularizes estimates of the means. We propose to use a flexible penalty that can be helpful for prioritizing genes that are differentially expressed in various ways. Here, we focus on a version of this penalty that enhances our ability to identify genes that are differentially expressed across viral status levels within each cell type. Note that if the *j*-th gene is differentially expressed across viral status levels within the *c*-th cell type, this implies *µ*_*jcv*_≠ *µ*_*jcv*_′ for some *v ≠ v*′. Our penalty encourages estimates of the means 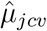 such that (*µ*_*jc*1_, …, *µ*_*jcV*_*)^⊤^ ∝ 1_*v*_* for many pairs (*j, c*). To accomplish this, we use the penalty

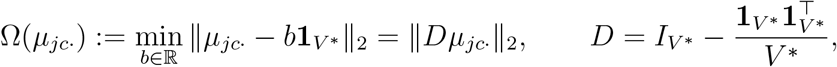

where *µ*_*jc*·_ = (*µ*_*jc*1_, …, *µ*_*jcV*_*)^⊤^. The full penalty is 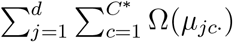 As can be seen from the first part of the definition above, this penalty using the Euclidean norm shrinks *µ*_*jc*·_ towards the nearest vector proportional to a vector of ones. As we just noted, if our estimate 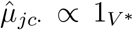, we would estimate that the mean expression of gene *j* in cell type *c* does not differ across viral statuses.

The penalty Ω was originally proposed in [31], and was recently used in [32]. In both works, the penalty was used to encourage “equi-sparsity” of regression coefficients in a distinct context. This penalty has never, to the best of our knowledge, been used in the context of mixture modeling. Typically, in mixture models, practitioners rely on pairwise fusion penalties (i.e., penalties like Σ*v*≠*v*′ |*µ*_*jcv*_ − *µ*_*jcv*_′|). Note that these penalties do not directly encourage 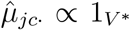, so they do not achieve the type of variable selection we seek. More-over, when *v*\* = 2, Ω is equivalent to pairwise fusion [32].

With this penalty in hand, we define our penalized maximum likelihood estimator of *θ*,

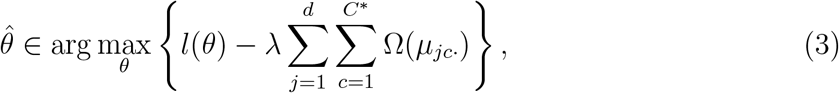

where *λ >* 0 is a user-specified tuning parameter (chosen by cross-validation). As *λ* → 0, the estimator tends towards the unpenalized maximum likelihood estimator. As *λ* → ∞, eventually the estimate 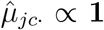 for all pairs (*j, c*). For intermediate values of *λ*, some 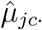 will be proportional to the vector of ones, whereas others will not.

The full details of the expectation-maximization algorithm that we use to estimate *θ* are provided in Additional file 1: Supplementary Methods. For this, we need to define the complete data log likelihood, i.e., the log-likelihood when (*C*_*i*_, v_*i*_) are observable for each *i* ∈ [*n*]. Let 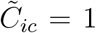 if *C*_*i*_ = *c* and zero otherwise, and likewise for 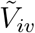. The complete data log-likelihood, ignoring constants not depending on *θ*, is

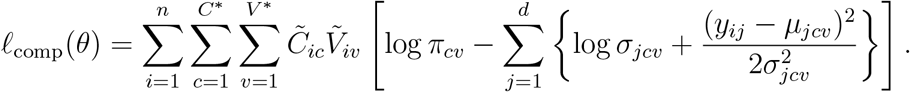

As mentioned, we select the tuning parameter *λ* using cross-validation. Specifically, we form *K* folds within each observed viral-status category and, separately, among cells with no observed viral-status label. For each fold *r* ∈ {1, …, *K*} and each candidate *λ* ∈ Λ, we fit the model on the training portion (the other *K* − 1 folds) and then evaluate the un-penalized data log-likelihood on the validation fold using the fitted parameters. We repeat this in parallel over folds, average the held-out log-likelihoods across *r*, and select the tuning parameter whose average log-likelihood is the largest. Details on the initialization and computations are provided in Additional file 1: Supplementary Methods.

### Permutation-based approximate inference procedure

After fitting the model to the generation set, we perform inference on the testing set without refitting the model. Let *n*_test_ denote the number of cells in the testing set. For each testing cell *k* ∈ [*n*_test_], we compute the *C*\* *× V*\* matrix

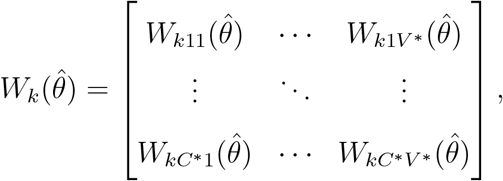

whose (*c, v*)-th entry 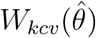 is the conditional probability that cell *k* has cell type *c* and viral status v, given its expression *Y*_*k*_ and whichever of *C*_*k*_ and v_*k*_ were observed, computed as if 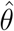 were the true parameter. When both *C*_*k*_ and v_*k*_ are observed, 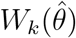 has a single nonzero entry equal to one. We refer to the 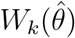 as weights. When a discrete label is needed, a cell is classified as infected if its weight on an infected state exceeds a user-specified threshold. With the weights in hand, we construct an approximate complete data log-likelihood for the testing cells,

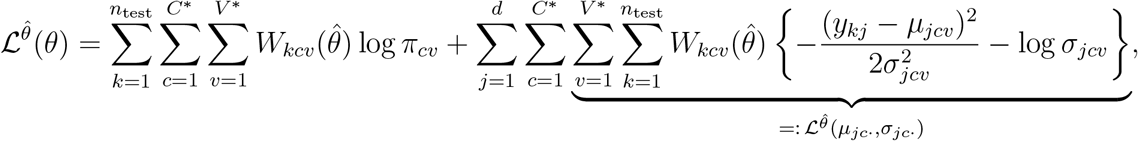

which is the complete data log-likelihood with the unobserved indicators 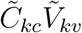 replaced by their conditional expectations under 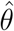. Two features of 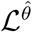 make it useful for differential expression testing. First, because 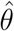 was estimated on the generation set, the weights do not depend on any parameter estimated from the testing cells. Second, apart from the mixing-proportion term, 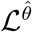 is a sum over genes and cell types of terms 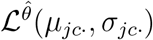, each depending only on the parameters for gene *j* in cell type *c*. Hence each of the *dC*\* gene and cell type combinations can be tested separately, using closed-form estimators, rather than by maximizing a constrained version of (1) for each hypothesis.

For gene *j* in cell type *c*, we test the null hypothesis 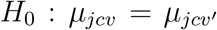 for all *v*≠ *v*′, that the mean expression is the same across viral statuses, against the alternative 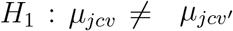 for some *v*≠ *v*′. The likelihood ratio statistic is

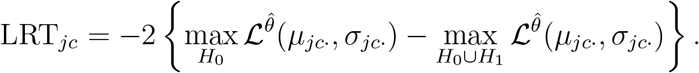

Under *H*_0_ ∪ *H*_1_, the maximizers are the weighted means and standard deviations

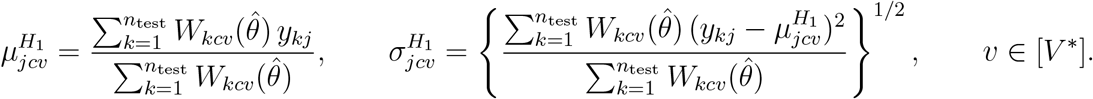

Under *H*_0_, the constrained maximizers of the mean and the variances would require an iterative algorithm to obtain. Instead, we approximate max 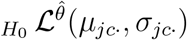 by plug-in estimate

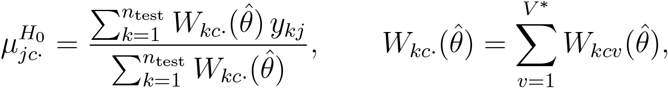

and setting 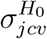 to be the same as the expression for 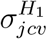 with 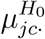 in place of 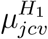. Our test statistic is thus

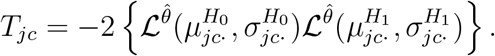

Instead of comparing *T*_*jc*_ to a reference distribution, as in standard likelihood ratio testing procedures, we rely on a permutation-based testing approach.

First, notice that 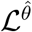 is an approximation to the complete data log-likelihood, not an observed data log-likelihood. There is then no reason, in general, that *T*_*jc*_ (or LRT_*jc*_) should have the usual asymptotic distribution. Second, although 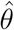 is independent of the testing set, the weight 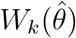 is not. The statistic LRT_*jc*_ thus inherits this information leakage. We therefore use a permutation reference distribution to obtain approximate *p*-values; exact validity requires the exchangeability conditions given below. Moreover, since *T*_*jc*_ is calibrated by permutation rather than by an asymptotic reference distribution, the use of plug-in estimators (rather than constrained maximum likelihood estimators) affects only the form of the statistic and not the validity of the test. For each testing cell *k* and cell type *c*, let

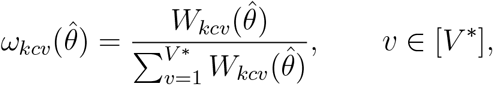

denote the estimated conditional probability that cell *k* has viral status *v* given that it has cell type *c*. Let *M* be the user-specified number of permutations (e.g., we take *M* = 10000 by default). Define *τ*_*m*_ as the *m*-th permutation of [*n*_test_], drawn uniformly within groups defined by the observed partitioning variable (e.g., sample or cell type). For example, for all cells of type *c, τ*_*m*_ randomly permutes cell indices. Next, define the permuted weights

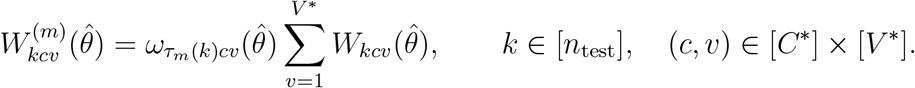

When permuting weights in this way, each cell retains its original cell type probabilities, 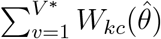, while its conditional viral status probabilities are exchanged with those of another cell in the same group.

If cell type is fully observed, this reduces to permuting the viral status probabilities among the cells within each cell type. Replacing 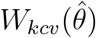 with 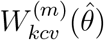 in the estimators above yields a null statistic 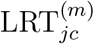, and the reported *p*-value for gene *j* in cell type *c* is

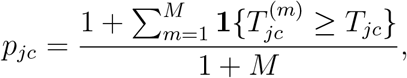

where **1**(·) is the indicator function. For genes not used to calculate the weights, the permutation p-value *p*_*jc*_ is valid if, conditional on the tested gene’s expression and the cell type weights, the conditional viral-status weight vectors are exchangeable within each permutation group. This condition may fail even when the tested gene is independent of the data used to calculate the weights, so we regard these p-values as approximate in practice.

The same *M* permutations are used for every gene, so for each *m* the null statistics for all genes are computed at once from weighted sums of *y*_*kj*_ and 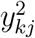. The cost of calibration is therefore *M* matrix products.

The genes used to estimate 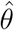 require a slightly more careful treatment. For such a gene *j*, the weight 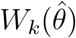 is a function of test set data *y*_*kj*_, so LRT_*jc*_ is computed with weights that were partly determined by the gene being tested. Accordingly, for each *m* ∈ [*M*], we permute 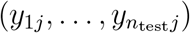 within each group while leaving all other genes fixed, recompute the weights for every testing cell from the permuted data with 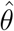 held fixed, and compute 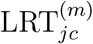 from the permuted expression and the recomputed weights. Because the log-weights are sums over the genes used to estimate 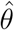, only the term for gene *j* changes under the permutation, so recomputing the weights requires one pass over the testing cells per permutation. For these genes, *p*_*jc*_ is valid if the tested gene’s expression is exchangeable within each permutation group, conditional on the other genes, observed labels, and fitted model. Neither exchangeability condition follows from the null hypothesis of equal means alone. Both conditions hold, for example, if cell type is observed, permutations stay within cell type, and, under *H*_0_, cells within each group are independent and identically distributed, with the tested gene independent of the other genes and observed viral-status labels. In practice, we treat the *p*-values for both sets of genes as approximate and assess their calibration in simulations.

To account for multiple testing, we use the Benjamini and Hochberg correction procedure. A user-defined significance level *α* is chosen, and genes with an adjusted *p*-value below *α* are considered differentially expressed between viral statuses within the given cell type.

## Data Description and Analysis

### Simulated data

Data was simulated to match the characteristics of scRNA-seq data using an extension to the scaffold R package [21]. Scaffold estimates parameters from a reference dataset and simulates data based on the experimental data generation process. We aimed to simulate data to explore scDEcrypter’s performance in data having infection heterogeneity at the cell type and sample levels. For all scenarios, we simulated data for three cell types, with cells evenly distributed among 498 infected cells and 1,497 uninfected cells. There were 19,969 total genes in the dataset, where gene-specific and cell-population level parameters were estimated from the influenza dataset [3] as a reference. The simulated data matched reference datasets in terms of sparsity and variability. To represent a wide variety of scenarios, we varied the proportion of cell type associated/marker genes, infection associated genes, and the fold-change (biological effect) of infection associated genes. There were 120 total simulation settings across all combinations of parameter settings shown in Table 1. For each setting, we generated 100 datasets. The fold-change of the cell type associated genes was fixed as two for all scenarios.

For each simulated dataset, scDEcrypter was run according to the scenario and pre-labeling parameter settings. All truly uninfected cells were left unlabeled (i.e. as NA). The default pre-processing, HvG selection procedure, and model fitting procedures in scDEcrypter were used with *C*\* = 3 and *v*\* = 2. For prediction, the threshold for infection state prediction was set to 0.5 and for the partial cell type labeling scenario, cell type prediction was set to the label with the largest probability. For inference, the LRT tested for differences between infected and uninfected cells within each cell type. We set *α* = 0.05 to determine significance on the adjusted p-values.

We used balanced accuracy to evaluate the performance of scDEcrypter. Performance for prediction was assessed by comparing each cell’s true (simulated) infection state label to the label assigned by scDEcrypter based on thresholding the weight matrix. Performance for differential expression was evaluated by comparing each gene’s true DE status to that determined by scDEcrypter using *α* = .05. Balanced accuracy accounts for both sensitivity and specificity, providing a more reliable measure of performance when class sizes are imbalanced. We also included the F1 score when comparing scDEcrypter to other methods.

### Computational benchmarking

We benchmarked scDEcrypter’s wall-clock runtime and peak memory usage on simulated datasets of varying size. We varied the number of cells (500–50,000), genes (100–10,000), and cell types (3, 12) one factor at a time, holding the others fixed. Each setting was replicated three times with random seeds. To simulate datasets of this size for computational benchmarking, we directly simulated counts as negative binomial with gene-wise means and dispersions estimated from the influenza reference dataset [3]. Cells were assigned uniformly at random to cell types and to a latent binary infection state, with 8% of genes carrying a cell type-associated effect and 5% an infection-associated effect (multiplicative effects drawn uniformly from [0.5, 2]). Five percent of cells, split evenly across infection states, were pre-labeled; the remainder were left unlabeled.

Each run followed the standard scDEcrypter pipeline: pre-processing (a 50/50 generation/test split stratified by label, variance stabilization, and HvG selection on the generation set), cross-validation of the penalty parameter, model fitting, and permutation-based DE testing on the held-out test set (10,000 permutations). Cross-validation and permutation testing were parallelized across eight CPU cores; all other steps were single-threaded.

### Influenza dataset

Data from Russell et al. [3] was obtained from the Gene Expression Omnibus (GSE108041). Cells from a human lung carcinoma cell-line were collected at 0, 6, 8, and 10 hours post infection, and the original study labeled 368 out of 12,447 cells as confidently infected using strict thresholds (about 3%). There were two replicates of the 8 hour time-point, and to avoid batch effects, we defined the additional partitioning variable as each of the distinct samples, so that *C*\* = 5 and *v*\* = 2. We pre-labeled cells with more than five influenza mRNA counts as infected, all 0-hour cells were labeled as uninfected, and all remaining cells with ambiguous or low-level viral expression were left unlabeled. The top HvGs were taken to be the unique collection of the 100 most highly variable genes in each virus-exposed sample. The threshold for exploring infection state prediction was set to 0.95. For inference, the permutation test tested for differences between infected and un-infected cell states at each time-point. Genes with an adjusted p-value less than 0.05 and an absolute logFC larger than 0.05 were considered DE.

Enrichment analysis on all DE genes was done using the `enrichPathway` function in the `ReactomePA` R package v1.48.0 [33] and visualized using the `enrichplot` R package v1.24.4 [34]. To identify increasing/decreasing genes, we subset those DE at either the 6-hr or 10-hr time-point and having an absolute logFC difference between the two time-points greater than 0.02 and having estimated 6-hr mean lower/higher than its 10-hr mean. To identify those decreasing over time and having a near return to uninfected baseline, we further filtered the decreasing genes to those with a 10-hr logFC lower than 0.20. Since some differences would be expected by 6-hr, we more leniently filtered the increasing trend genes to those having a logFC less than 0.20 at 6-hr. Enrichment of DE genes with trends over time was done using the enrichR package v3.4 using the ‘GO Biological Process 2026’ and ‘Reactome Pathways 2024’ databases.

### SAR-Cov-2 dataset

Data from Ravindra et al. [14] was obtained from the Gene Expression Omnibus (accession GSE166766). Human bronchial epithelial cells were experimentally infected with SARS-Cov-2 and collected at 1, 2, and 3 days post-infection and a control sample. The original study defined cells as infected based on having at least ten viral read counts. Based on the original annotations, we set cell type as the additional partitioning variable and set *C*\* = 8. In this analysis, we set *v*\* = 3 to explore whether scDEcrypter could distinguish between infected and bystander states. To mitigate batch effects, scDEcrypter was run independently and identically for each day post-infection along with the uninfected control cells. We pre-labeled cells with more than 40 viral mRNA counts as infected and uninfected cells having zero viral reads as uninfected, with all remaining cells left unlabeled. To allow additional information sharing between cell types in the estimation stage, we pre-labeled only 75% of cells, randomly. The top HvGs were taken to be the unique collection of the 120 most highly variable genes from each cell type in the exposed sample, as well as, 100 additional HvGs across cell types in the uninfected control sample to assist with discriminating cell type membership for unlabeled cells. The threshold for exploring infection state was set to 0.5. For the exploratory cell type prediction with partially labeled test set data, the cell type prediction threshold was also set to 0.5. For inference, the permutation test tested the unique collection of the top 2,000 genes from each cell type for differences between infected versus bystander states and bystander versus uninfected states. For all comparisons and cell types, genes with adjusted p-value less than 0.05 and absolute logFC larger than 1 were considered DE. Enrichment of DE genes was done using the enrichR package v3.4 using the ‘GO Biological Process 2026’ and ‘Reactome Pathways 2024’ databases.

### Existing approaches

We compared scDEcrypter to existing methods used in scRNA-seq DE analysis: Seurat [35], DESeq2 [17], and MAST [16]. For the simulation comparisons, the cell group labels were set to mimic workflows used in practice where cells labeled as infected matched those used in the pre-labeling for scDEcrypter and all other cells were labeled as uninfected. For Seurat, we used the default Wilcoxon rank-sum test implementation with pre-testing filtering turned off (i.e., min.pct=0 and logfc.threshold=0). For DESeq2 and MAST, the default settings were used.

We also compared scDEcrypter to scANvI [18], a semi-supervised deep generative model that learns a latent representation with a variational autoencoder and incorporates a classifier trained on labeled cells to predict cell states for unlabeled cells. Because scANvI requires labeled examples of every state of interest, 5% of infected and uninfected cells within each cell type were pre-labeled (overall 10% pre-labeling), with all remaining training cells treated as unlabeled. scANvI was evaluated using the same simulation scenarios and matched train-test split datasets. Highly variable genes and model parameters were learned using training cells only; infection-state prediction and DE detection were then evaluated in held-out test cells. We compared scANvI to scDEcrypter’s performance at 10% pre-labeled cells.

We used a fixed scvi-tools configuration across simulations, without dataset-specific hyperparameter tuning. SCvI was trained for 100 epochs with a 30-dimensional latent space and two hidden layers. SCANvI was initialized from the fitted SCvI model and trained for 25 epochs using a two-layer classifier with dropout rate 0.1. Because native scANvI DE is restricted to genes used for model fitting, DE was assessed using Wilcoxon rank-sum tests between scANvI-predicted infection groups in the held-out test set, with Benjamini– Hochberg correction as implemented in Scanpy. Up to 5,000 highly variable genes, selected from the training cells, were included in DE testing.

## Declarations

### Ethics approval and consent to participate

Not applicable.

### Consent for publication

Not applicable.

### Availability of data and materials

The scDEcrypter package is available at https://github.com/bachergroup/scDEcrypter. A snapshot of the release of scDEcrytper v1.0.0 used in this manuscript has been archived at Zenodo (https://doi.org/10.5281/zenodo.23093820). We obtained the influenza and SARS-Cov-2 datasets from Gene Expression Omnibus (GEO) with accession numbers GSE108041 [3] and GSE166766 [14].

### Competing interests

The authors declare no competing interests.

### Funding

The project was supported by the National Institutes of Health under Award Numbers: R35GM146895, P01CA214091, and R01AI108407.

### Authors’ contributions

LZ implemented the software, designed and performed simulation studies, and assisted with pre-processing and analyzing the scRNA-seq datasets. KE and ST assisted with biological interpretation of case-study results. AM designed the scDEcrypter method, provided critical input on theoretical formulation, and assisted in writing codes for the penalized mixture modeling framework, inference strategy, and simulation studies. RB conceived the study, designed the scDEcrypter method, contributed to the statistical and biological interpretations, analyzed the scRNA-seq datasets, and supervised the project. All authors contributed to manuscript writing, revising, and figure preparation. All authors discussed the results, contributed to the final manuscript, and approved its submission.

## Acknowledgments

Portions of the R code were translated to C++ with assistance from the large language model Claude Fable 5 (Anthropic) to improve computational performance. All generated code was reviewed, tested, and validated by the authors. We thank student Rebecca Lincoln from the University of Florida for her critical review of the manuscript.

## Notes

### Competing Interest Statement

The authors have declared no competing interest.

### Summary of Updates

Implemented permutation approach to obtain p-values for DE testing, additional simulation studies and benchmarking, code/package acceleration.

## References

[1] Jones JE, Le Sage V, Lakdawala SS. Viral and host heterogeneity and their effects on the viral life cycle. Nature Reviews Microbiology. 2021 Apr;19(4):272–82.

[2] Steuerman Y, Cohen M, Peshes-Yaloz N, Valadarsky L, Cohn O, David E, et al. Dissection of Influenza Infection In Vivo by Single-Cell RNA Sequencing. Cell Systems. 2018 Jun;6(6):679-91.e4.

[3] Russell AB, Trapnell C, Bloom JD. Extreme heterogeneity of influenza virus infection in single cells. Elife. 2018;7:e32303.

[4] Banerjee A, El-Sayes N, Budylowski P, Jacob RA, Richard D, Maan H, et al. Experimental and natural evidence of SARS-CoV-2-infection-induced activation of type I interferon responses. iScience. 2021 May;24(5):102477.

[5] Song E, Zhang C, Israelow B, Lu-Culligan A, Prado AV, Skriabine S, et al. Neuroinvasion of SARS-CoV-2 in human and mouse brain. Journal of Experimental Medicine. 2021 Mar;218(3):e20202135.

[6] Douam F, Hrebikova G, Albrecht YES, Sellau J, Sharon Y, Ding Q, et al. Singlecell tracking of flavivirus RNA uncovers species-specific interactions with the immune system dictating disease outcome. Nature Communications. 2017 Mar;8(1):14781.

[7] Waickman AT, Friberg H, Gromowski GD, Rutvisuttinunt W, Li T, Siegfried H, et al. Temporally integrated single cell RNA sequencing analysis of PBMC from experimental and natural primary human DENV-1 infections. PLoS pathogens. 2021 Jan;17(1):e1009240.

[8] Moore KM, Pelletier AN, Lapp S, Metz A, Tharp GK, Lee M, et al. Single-cell analysis reveals an antiviral network that controls Zika virus infection in human dendritic cells. Journal of Virology. 2024 May;98(5):e00194–24.

[9] Swaminath S, Russell AB. The use of single-cell RNA-seq to study heterogeneity at varying levels of virus–host interactions. PLOS Pathogens. 2024 Jan;20(1):e1011898.

[10] Whitmore LS, Tisoncik-Go J, Gale M. scPathoQuant: a tool for efficient alignment and quantification of pathogen sequence reads from 10× single cell sequencing datasets. Bioinformatics. 2024 Mar;40(4):btae145.

[11] Cohen P, DeGrace EJ, Danziger O, Patel RS, Barrall EA, Bobrowski T, et al. Unambiguous detection of SARS-CoV-2 subgenomic mRNAs with single-cell RNA sequencing. Microbiology Spectrum. 2023 Oct;11(5):e00776–23.

[12] Sanjuán R, Domingo-Calap P. Mechanisms of viral mutation. Cell Mol Life Sci. 2016 Dec;73(23):4433–48.

[13] Lun ATL, McCarthy DJ, Marioni JC. A step-by-step workflow for low-level analysis of single-cell RNA-seq data with Bioconductor. F1000Research. 2016 Oct;5:2122.

[14] Ravindra NG, Alfajaro MM, Gasque V, Huston NC, Wan H, Szigeti-Buck K, et al. Single-cell longitudinal analysis of SARS-CoV-2 infection in human airway epithelium identifies target cells, alterations in gene expression, and cell state changes. PLOS Biology. 2021 Mar;19(3):e3001143.

[15] Butler A, Hoffman P, Smibert P, Papalexi E, Satija R. Integrating single-cell transcriptomic data across different conditions, technologies, and species. Nature biotechnology. 2018;36(5):411–20.

[16] Finak G, McDavid A, Yajima M, Deng J, Gersuk V, Shalek AK, et al. MAST: a flexible statistical framework for assessing transcriptional changes and characterizing heterogeneity in single-cell RNA sequencing data. Genome biology. 2015;16:1–13.

[17] Love MI, Huber W, Anders S. Moderated estimation of fold change and dispersion for RNA-seq data with DESeq2. Genome biology. 2014;15:1–21.

[18] Xu C, Lopez R, Mehlman E, Regier J, Jordan MI, Yosef N. Probabilistic harmonization and annotation of single-cell transcriptomics data with deep generative models. Molecular systems biology. 2021;17(1):e9620.

[19] Missarova A, Dann E, Rosen L, Satija R, Marioni J. Leveraging neighborhood representations of single-cell data to achieve sensitive DE testing with miloDE. Genome biology. 2024;25(1):189.

[20] Madrigal A, Lu T, Soto LM, Najafabadi HS. A unified model for interpretable latent embedding of multi-sample, multi-condition single-cell data. Nature Communications. 2024;15(1):6573.

[21] Bacher R, Chu LF, Argus C, Bolin JM, Knight P, Thomson JA, et al. Enhancing biological signals and detection rates in single-cell RNA-seq experiments with cDNA library equalization. Nucleic Acids Research. 2022;50(2):e12–2.

[22] Watanabe T, Watanabe S, Kawaoka Y. Cellular networks involved in the influenza virus life cycle. Cell host & microbe. 2010;7(6):427–39.

[23] Long JS, Giotis ES, Moncorgé O, Frise R, Mistry B, James J, et al. Species difference in ANP32A underlies influenza A virus polymerase host restriction. Nature. 2016 Jan;529(7584):101–4.

[24] Staller E, Sheppard CM, Neasham PJ, Mistry B, Peacock TP, Goldhill DH, et al. ANP32 Proteins Are Essential for Influenza Virus Replication in Human Cells. Journal of Virology. 2019 Sep;93(17):e00217–9.

[25] Chen G, Liu CH, Zhou L, Krug R. Cellular DDX21 RNA Helicase Inhibits Influenza A Virus Replication but Is Counteracted by the Viral NS1 Protein. Cell Host & Microbe. 2014 Apr;15(4):484–93.

[26] Meng X, Zhu Y, Yang W, Zhang J, Jin W, Tian R, et al. HIF-1α promotes virus replication and cytokine storm in H1N1 virus-induced severe pneumonia through cellular metabolic reprogramming. Virologica Sinica. 2024 Feb;39(1):81–96.

[27] Gupta P, Sahu R, Nandi I, Setty SRG, Das S, Singh M. Interaction and co-phase separation of SARS-CoV-2 nucleocapsid protein and human hnRNPA1 and its implications for viral life cycle. FEBS Letters. 2026 May:1873-3468.70375.

[28] Zhang H, Wang Z, Nguyen HTT, Watson AJ, Lao Q, Li A, et al. Integrin α5 β1 contributes to cell fusion and inflammation mediated by SARS-CoV-2 spike via RGD-independent interaction. Proceedings of the National Academy of Sciences. 2023 Dec;120(50):e2311913120.

[29] Tugaeva KV, Sysoev AA, Kapitonova AA, Smith JLR, Zhu P, Cooley RB, et al. Human 14-3-3 Proteins Site-selectively Bind the Mutational Hotspot Region of SARS-CoV-2 Nucleoprotein Modulating its Phosphoregulation. Journal of Molecular Biology. 2023 Jan;435(2):167891.

[30] Ahlmann-Eltze C, Huber W. Comparison of transformations for single-cell RNA-seq data. Nature Methods. 2023;20(5):665–72.

[31] Molstad AJ, Motwani K. Multiresolution categorical regression for interpretable celltype annotation. Biometrics. 2023;79(4):3485–96.

[32] Fu J, Molstad AJ, Zou H. A direct approach to tree-guided feature aggregation for high-dimensional regression. arXiv preprint arXiv:250719650. 2025.

[33] Yu G, He QY. ReactomePA: an R/Bioconductor package for reactome pathway analysis and visualization. Molecular BioSystems. 2016;12(2):477–9.

[34] Yu G. enrichplot: Visualization of Functional Enrichment Result; 2025. R package version 1.24.4.

[35] Satija R, Farrell JA, Gennert D, Schier AF, Regev A. Spatial reconstruction of single-cell gene expression data. Nature biotechnology. 2015;33(5):495–502.

