## Additional File 1 for "scDEcrypter: Uncertainty-aware differential expression analysis for viral infection in scRNA-seq"

**1 Supplementary Methods** **2**

**3 Supplementary Figures** **19**

### 1 Supplementary Methods

#### 1.1 Expectation-maximization algorithm

The expectation (E) step computes the conditional expectation of the complete-data log-likelihood given the observed data and current parameter iterate. The subsequent block-wise updates increase this expected log-likelihood minus the penalty. Let $l(\theta)$ denote the observed-data log-likelihood and let $D = I_{V^*} - \mathbf{1}_{V^*}\mathbf{1}_{V^*}^\top/V^*$. The penalty is $\Omega(\mu_{jc\cdot}) = \|D\mu_{jc\cdot}\|_2$.

$$\hat{\theta} \in \arg\max_{\theta} \left\{ l(\theta) - \lambda \sum_{j=1}^{d} \sum_{c=1}^{C^*} \Omega(\mu_{jc\cdot}) \right\}, \tag{1}$$

The parameter space restricts $\sigma^2_{jcv} \geq 0.1$ for all $(j, c, v)$. The software's tuning input $\lambda_0$ is a per-cell penalty value. For a fit with $n$ cells, the coefficient in (1) is $\lambda = n\lambda_0$; a cross-validation training fold with $n_{\text{train}}$ cells uses $\lambda = n_{\text{train}}\lambda_0$.

##### 1.1.1 Expectation step

As in the main text, let $\tilde{C}_{ic}$ and $\tilde{V}_{iv}$ indicate whether cell $i$ has cell type $c$ and viral status $v$, respectively. Up to a constant independent of $\theta$, the complete-data log-likelihood is

$$\ell_{\text{comp}}(\theta) = \sum_{i=1}^{n} \sum_{c=1}^{C^*} \sum_{v=1}^{V^*} \tilde{C}_{ic}\tilde{V}_{iv} \left\{ \log \pi_{cv} + \sum_{j=1}^{d} \left[ -\frac{(y_{ij} - \mu_{jcv})^2}{2\sigma^2_{jcv}} - \log \sigma_{jcv} \right] \right\}.$$

At iteration $k$, the E-step replaces each unobserved product $\tilde{C}_{ic}\tilde{V}_{iv}$ by its conditional expectation $W^{(k)}_{icv}$ under $\theta^{(k)}$, given the observed data. In the following, let $Y_i = (Y_{i1}, \ldots, Y_{id})^\top$ and similarly for $y_i$. For each cell $i$, let $\mathcal{A}_i$ contain the $(c, v)$ pairs compatible with its observed cell-type and viral-status labels. Using the index sets $\mathcal{B}$, $\mathcal{C}$, $\mathcal{V}$, and $\mathcal{N}$ defined in the main text,

$$\mathcal{A}_i = \begin{cases} \{(c_i, v_i)\}, & i \in \mathcal{B}, \\ \{(c_i, v) : v \in [V^*]\}, & i \in \mathcal{C}, \\ \{(c, v_i) : c \in [C^*]\}, & i \in \mathcal{V}, \\ [C^*] \times [V^*], & i \in \mathcal{N}. \end{cases}$$

All four patterns of observed and unobserved labels are covered by

$$W_{icv}^{(k)} = \frac{\mathbb{1}\{(c, v) \in \mathcal{A}_i\}\, \pi_{cv}^{(k)} \prod_{j=1}^{d} f_{ij}^{\theta^{(k)}}(y_{ij} \mid c, v)}{\sum_{(c',v') \in \mathcal{A}_i} \pi_{c'v'}^{(k)} \prod_{j=1}^{d} f_{ij}^{\theta^{(k)}}(y_{ij} \mid c', v')}.$$

Thus, states that conflict with an observed label receive weight zero. When both labels are observed, the observed pair receives weight one.

###### 1.1.2 Maximization step

Given the E-step weights, the generalized M-step updates $\theta$ to increase the penalized $Q$ function:

$$\begin{aligned} Q_\lambda(\theta \mid \theta^{(k)}) &= \mathrm{E}_{\theta^{(k)}}[\ell_{\mathrm{comp}}(\theta) \mid \text{observed data}] - \lambda \sum_{j=1}^{d} \sum_{c=1}^{C^*} \Omega(\mu_{jc\cdot}) \\ &= \sum_{i=1}^{n} \sum_{c=1}^{C^*} \sum_{v=1}^{V^*} W_{icv}^{(k)} \left\{ \log \pi_{cv} + \sum_{j=1}^{d} \left[ -\frac{(y_{ij} - \mu_{jcv})^2}{2\sigma_{jcv}^2} - \log \sigma_{jcv} \right] \right\} \\ &\quad - \lambda \sum_{j=1}^{d} \sum_{c=1}^{C^*} \Omega(\mu_{jc\cdot}). \end{aligned}$$

Instead of maximizing the $Q$ function with respect to $\theta$, we find the $(k+1)^{\text{th}}$ iterate value, $\theta^{(k+1)}$ such that

$$Q_\lambda(\theta^{(k+1)} | \theta^{(k)}) \geq Q_\lambda(\theta^{(k)} | \theta^{(k)}).$$

This is a generalized expectation-maximization (GEM) algorithm with blockwise updates.

We update the $\mu_{jcv}$ with the $\sigma_{jcv}$ fixed at their previous iterate, then update the $\sigma_{jcv}$ with $\mu_{jcv}$ fixed at the new iterate. The $\pi_{cv}$ can be updated independently of the means and variances using

$$\pi_{cv}^{(k+1)} = \frac{\sum_{i=1}^{n} W_{icv}^{(k)}}{n}. \tag{2}$$

Then, to update the $\mu_{jcv}$ with the $\sigma_{jcv}$ fixed at $\sigma_{jcv}^{(k)}$, we need to maximize

$$\left\{\sum_{i=1}^{n}\sum_{j=1}^{d}\sum_{c=1}^{C^*}\sum_{v=1}^{V^*} W_{icv}^{(k)}\left[-\frac{(y_{ij}-\mu_{jcv})^2}{2(\sigma_{jcv}^{(k)})^2}\right]\right\} - \lambda\sum_{j=1}^{d}\sum_{c=1}^{C^*}\Omega(\mu_{jc\cdot}).$$

The above function is separable across the $d$ genes and $C^*$ cell types, so we need to only concern ourselves with maximizing

$$\left\{\sum_{i=1}^{n}\sum_{v=1}^{V^*} W_{icv}^{(k)}\left[-\frac{(y_{ij}-\mu_{jcv})^2}{2(\sigma_{jcv}^{(k)})^2}\right]\right\} - \lambda\Omega(\mu_{jc\cdot}),$$

separately for each $(j, c) \in [d] \times [C^*]$. When all viral-status weight totals for $(j, c)$ are positive, this problem can be expressed as

$$\underset{m\in\mathbb{R}^{V^*}}{\text{minimize}} \quad \frac{1}{2}\|S^{-1/2}(m-\bar{y}_{jc})\|_2^2 + \lambda\|Dm\|_2 \tag{3}$$

where $S = \text{Diag}(\frac{(\sigma_{jc1}^{(k)})^2}{\sum_{i=1}^{n} W_{ic1}^{(k)}}, ..., \frac{(\sigma_{jcV^*}^{(k)})^2}{\sum_{i=1}^{n} W_{icV^*}^{(k)}})$ and $\bar{y}_{jc} = (\bar{y}_{jc1}, \ldots, \bar{y}_{jcV^*})^\top$ with

$$\bar{y}_{jcv} = \frac{\sum_{i=1}^{n} y_{ij} W_{icv}^{(k)}}{\sum_{i=1}^{n} W_{icv}^{(k)}}, \quad v \in [V^*].$$

The problem (3) is a convex optimization problem and can be solved using the proximal gradient descent algorithm. The proximal gradient descent algorithm is primarily used to solve non-smooth convex optimization problems that are separable into a smooth and a non-smooth term. In our case, the smooth term is $f(m) = \frac{1}{2}\|S^{-1/2}(m-\bar{y}_{jc})\|_2^2$ and non-smooth term is $\lambda\|Dm\|_2$. The $(s+1)^{\text{th}}$ iterate of the algorithm to solve (3) is given by

$$m^{(s+1)} = \arg\min_m \left\{ \frac{1}{2}\|m - \{m^{(s)} - \alpha\nabla f(m^{(s)})\}\|_2^2 + \alpha\lambda\|Dm\|_2 \right\},$$

The software starts at $m^{(0)} = \mu_{jc\cdot}^{(k)}$ and uses the step size $\alpha = 1/\max_v\{\sum_i W_{icv}^{(k)}/(\sigma_{jcv}^{(k)})^2\}$. This choice ensures that each proximal-gradient step does not increase the objective in (3). The final proximal-gradient iterate becomes $\mu_{jc\cdot}^{(k+1)}$. If any viral-status weight total is zero, the software retains the current vector $\mu_{jc\cdot}^{(k)}$. When $\lambda = 0$, the update reduces to a gradient descent step. The proximal update has a closed form for any $V^*$; see Molstad and Motwani [2023] or Fu et al. [2025].

Using the resulting $\mu_{jc\cdot}^{(k+1)}$, we update $\sigma_{jcv}^2$ by

$$(\sigma_{jcv}^2)^{(k+1)} = \max\left\{0.1, \frac{\sum_{i=1}^n W_{icv}^{(k)}(y_{ij} - \mu_{jcv}^{(k+1)})^2}{\sum_{i=1}^n W_{icv}^{(k)}}\right\}. \tag{4}$$

If $\sum_i W_{icv}^{(k)} = 0$, the implementation sets $(\sigma_{jcv}^2)^{(k+1)} = 0.1$.

As in the main text, the iterate collects the means, variances, and mixing proportions in that order:

$$\begin{aligned}\theta^{(k)} = \big(&\mu_{111}^{(k)}, \dots, \mu_{dC^*V^*}^{(k)},\\ &(\sigma_{111}^2)^{(k)}, \dots, (\sigma_{dC^*V^*}^2)^{(k)},\\ &\pi_{11}^{(k)}, \dots, \pi_{C^*V^*}^{(k)}\big)^\top.\end{aligned}$$

Within each GEM iteration, we update the mixing proportions by (2), the means using (3), and then the variances by (4). Each block update maximizes or increases $Q_\lambda$ with the other blocks fixed. Hence $Q_\lambda(\theta^{(k+1)} \mid \theta^{(k)}) \geq Q_\lambda(\theta^{(k)} \mid \theta^{(k)})$, which gives the ascent property for the penalized observed-data log-likelihood.

#### 1.2 Cross-validation initialization

Each training-fold fit starts with uniform mixing proportions, $\pi_{cv}^{(0)} = 1/(C^*V^*)$, variances $(\sigma_{jcv}^2)^{(0)} = 1$, and a data-driven mean array $M^{(0)} = (\mu_{jcv}^{(0)})$. For each gene and $(c, v)$ pair, the initial mean uses cells with both labels observed. If no such cells are available, the code uses, in order, the mean for cell type $c$, the mean for viral status $v$, and the overall gene mean:

$$M_{jcv}^{(0)} \leftarrow \begin{cases} \text{mean}\big(Y_{ij} : C_i = c,\, V_i = v\big), & \text{if available,} \\ \text{mean}\big(Y_{ij} : C_i = c\big), & \text{else if available,} \\ \text{mean}\big(Y_{ij} : V_i = v\big), & \text{else if available,} \\ \text{mean}\big(Y_{ij}\big), & \text{otherwise.} \end{cases}$$

Starting from these values, the code first runs an unpenalized EM fit. Every candidate penalty starts from this same fitted initialization. The selected penalty maximizes the average observed-data log-likelihood on the validation folds.

#### 1.3 Alternative penalty variation

In our implementation, we apply the penalty to the rows of the mean matrices $\mu_{jc\cdot}$, thus encouraging estimates that have many gene and cell type combinations wherein the mean expression does not differ across viral status levels. In some contexts, alternative penalization schemes may be more appropriate. For example, if one assumed that for many genes, mean expression did not differ across either cell types or viral status, we would recommend using a version of the penalty given by

$$\sum_{j=1}^{d} \Omega_{\text{all}}(\mu_{j\cdot\cdot}) = \sum_{j=1}^{d} \min_{a \in \mathbb{R}} \|\mu_{j\cdot\cdot} - a\mathbf{1}_{C^*}\mathbf{1}_{V^*}^\top\|_F = \sum_{j=1}^{d} \|D_{C^*V^*}\text{vec}(\mu_{j\cdot\cdot})\|_2,$$

where $D_{C^*V^*} = I_{C^*V^*} - \mathbf{1}_{C^*V^*}\mathbf{1}_{C^*V^*}^\top/(C^*V^*)$, vec stacks the columns of its matrix-valued argument, and $\|\cdot\|_F$ is the Frobenius norm. One could also combine the two penalties, assigning a distinct tuning parameter for each. For example, a natural estimator is

$$\tilde{\theta} \in \arg\max_{\theta} \left\{ l(\theta) - \lambda_1 \sum_{j=1}^{d} \sum_{c=1}^{C^*} \Omega(\mu_{jc\cdot}) - \lambda_2 \sum_{j=1}^{d} \Omega_{\text{all}}(\mu_{j\cdot\cdot}) \right\}. \tag{5}$$

We use (1) in our implementation in order to limit the number of tuning parameters that need to be chosen.

### 2 Simulation studies under model misspecification

To assess various aspects of scDEcrypter's performance under model misspecification, we performed additional simulation studies. In particular, we consider two new data generating models. In the first model, where the data are generated to be count-valued, we study how non-normality, varying infection intensity, correlation from shared latent factors, and cell-specific library size affect our method. In the second model, we generate data from a multivariate normal so that expression is correlated in an interpretable way. In each setting below, we perform 50 independent replications. In all settings, we generate datasets with 1,995 cells, three equally sized cell types, 166 infected cells per type, 19,969 non-viral genes, and 8 viral genes. All analyses use a 50:50 generation/test split, five-fold penalty selection, and 1000 test-set permutations. Tables report replication means with Monte Carlo standard errors in parentheses.

#### 2.1 Data generating models

##### 2.1.1 Negative binomial expression counts (Model 1)

Our main simulations used Scaffold with parameters estimated from the influenza data. For this study, we used the simplified negative-binomial model below, with parameters chosen to produce a comparable sequencing depth to the main studies. For Model 1, we model the counts for host gene $j$ in cell $i$ as

$$Y_{ij} \mid \mu_{ij}, \phi_j \sim \mathrm{NB}(\mu_{ij}, \phi_j), \qquad \phi_j = 0.3 + \exp(-A_j), \tag{6}$$

where $\mathrm{NB}(\mu, \phi)$ has mean $\mu$ and variance $\mu + \phi\mu^2$. Conditional on the variables described below, the host counts are independent across cells and genes and their means satisfy

$$\log(\mu_{ij}) = A_j + U_i + (\log 2)\mathbf{1}\{j \text{ marker gene for cell type } c_i\} + \xi_i\beta_j + \sum_{k=1}^{3} L_{jk}F_{ik}. \tag{7}$$

The terms appearing in (7) are generated as follows.

- *Baseline expression.* Each host gene has $A_j \sim \mathrm{N}(-2.5, 1.6^2)$.
- *Library size.* $U_i \sim \mathrm{N}(0, \sigma_U^2)$ scales every host gene in cell $i$ by $s_i = \exp(U_i)$. By default, we take $\sigma_U = 0.35$.
- *Cell-type markers.* The indicator $\mathbf{1}\{j \text{ marker gene for cell type } c_i\}$ doubles a marker gene's mean in its assigned type. For each replication, 998 genes are randomly selected as cell-type markers, and each one is assigned to one cell type.
- *Infection-associated genes.* A set of 998 genes, disjoint from the marker genes is randomly selected to have an infection effect. For each infection-associated gene, we set $\beta_j = S_j u_j \log f$, where $\Pr(S_j = -1) = \Pr(S_j = 1) = 1/2$ and $u_j \sim \mathrm{Unif}(0.5, 1.5)$ indicates the signs and magnitudes, respectively. For all other genes, $\beta_j = 0$.
- *Infection intensity.* For each cell, infection intensity is $\xi_i = Z_i B_i$, with $B_i \sim \mathrm{Beta}(a, 1)$ (with $a = 2$ being the default) and $Z_i = 1$ for an infected cell and $Z_i = 0$ otherwise. We refer to $a$, which controls the distribution of $B_i$ as the infection intensity parameter. Uninfected cells have $\xi = 0$. In an infected cell, the infection-associated genes are scaled by $B_i$. Thus, for an infection-associated gene with $u_j = 1$ in a cell with full infection intensity $B_i = 1$, the mean fold change is $f$ for an upregulated gene and $1/f$

for a downregulated gene. Other values of $u_j$ and $B_i$ change the magnitude of this effect.

- *Shared factors.* Each cell has three scores $(F_{i1}, F_{i2}, F_{i3})$ where independently, $F_{ik} \sim \mathrm{N}(0, 1)$. For each factor $k$, 10% of genes are randomly selected (without replacement) and given independent loadings $L_{jk} \sim \mathrm{N}(0, \sigma_L^2)$. All other loadings are set to zero. As a default, we set $\sigma_L = 0.35$. Thus, the sum $\sum_k L_{jk}F_{ik}$ creates dependence among genes that share a cell-level factor.

Except for the fixed-size selections of infected cells and gene sets, all random draws above are independent.

Under this construction, the difference in gene means between infected and uninfected cells can be characterized through

$$\log(\mu_{ij}) = \begin{cases} A_j + U_i + (\log 2)\mathbf{1}\{j \text{ marks } c_i\} + \sum_{k=1}^{3} L_{jk}F_{ik}, & Z_i = 0, \\ A_j + U_i + (\log 2)\mathbf{1}\{j \text{ marks } c_i\} + B_i\beta_j + \sum_{k=1}^{3} L_{jk}F_{ik}, & Z_i = 1. \end{cases}$$

An uninfected cell has no infection term. An infected cell has the gene-specific effect $\beta_j$ scaled by its intensity $B_i$. For genes outside the infection-associated set, $\beta_j = 0$ in both cases.

To generate viral reads for viral genes, let $(r_1, \ldots, r_8) = (0.30, 0.20, 0.15, 0.10, 0.10, 0.06, 0.05, 0.04)$. After fixing each cell's infection status, infection intensity, library size, and the other variables that determine the count means, we draw the viral segment counts independently of one another and of the host counts, with

$$R_{ih} \sim \mathrm{NB}(r_h\{100 s_i \xi_i^3 e^{E_i} + 0.05\}, 0.5), \qquad h = 1, \ldots, 8,$$

where $E_i \sim \mathrm{N}(0, 0.5^2)$, the viral count noise, is drawn independently of all other variables. For uninfected cells, $\xi_i = 0$ and the mean viral count is only $0.05 r_h$, representing the potential for low levels of viral reads as observed in scRNA-seq datasets. For infected cells, the mean also contains $100 r_h s_i B_i^3 e^{E_i}$, allowing viral read means to increase with infection intensity.

We evaluate pre-labeling 1,995$*q$ cells in two ways: either uniformly among infected cells or by choosing those with the largest total viral count $\sum_h R_{ih}$. We denote uniform pre-labeling by setting $\pi_V = 0$, and viral count-based pre-labeling by $\pi_V = 1$ in the tables below. The same scDEcrypter pipeline (with processing and normalization) was carried out for all simulations: a 50:50 generation/test split, five-fold selection of the tuning parameter, and permutation-based DE testing on the test set with 1000 permutations.

##### 2.1.2 Correlated multivariate normal (Model 2)

Model 2 uses the same cell and gene assignment rules as Model 1. Here $f \in \{1.5, 2, 4\}$, infection intensity is binary ($\xi_i = Z_i$), and pre-labels are selected uniformly among infected cells. Expression is simulated as:

$$X_{ij} = (\log 2)\mathbf{1}\{j \text{ marks cell type } c_i\} + Z_i\beta_j + \epsilon_{ij},$$

where the vectors $\epsilon_i = (\epsilon_{i1}, \dots, \epsilon_{id})^\top \sim \mathrm{N}_d(0, \Sigma)$ are independent across cells, with $\Sigma_{j,k} = \rho^{|j-k|}$. In the simulations below, we consider $\rho \in \{0.3, 0.5, 0.7, 0.9, 0.95, 0.99, 0.999\}$. Although Model 1 also induces gene-gene correlation, the correlation induced by Model 2 is more straightforward and interpretable. scDEcrytper was run with the standard workflow excluding variance stabilization.

#### 2.2 Simulation study designs and results

##### 2.2.1 Viral-load-dependent pre-labeling

Host and viral counts were first generated following the negative-binomial model detailed above. Under Model 1, infection intensity in infected cells follows a Beta(2, 1), and three sparse cell-level factors induce host-gene dependence. We compared two pre-labeling strategies: uniform sampling of truly infected cells and selecting cells having the highest total viral counts. Figure R1 displays results under configurations of $f \in \{1.5, 2, 4\}$ and pre-label proportions $q \in \{0.01, 0.04, 0.10\}$. Table R1 gives the $f = 2$ results. Infection balanced accuracy averages the proportion of infected and uninfected unlabeled test cells classified correctly. DE F1 is the harmonic mean of precision and recall for truly DE gene by cell-type pairs.

In Figure R1, we see that viral-based pre-labeling favors cells with high infection intensity, perhaps unsurprisingly. Mean intensity among pre-labeled cells was 0.92 at $q = 0.01$ and 0.85 at $q = 0.10$, versus 0.67 among all infected cells. Infection balanced accuracy remained 0.84–0.87 with viral-load labels, but DE F1 was below 0.06 at $f = 2$. We also report the fraction of tested null gene–cell-type pairs with raw p-values at or below 0.05. At $f = 2$ and $q = 0.04$, this fraction is 0.165 under random pre-labeling and 0.186 under viral-load pre-labeling. The corresponding fractions called DE after BH adjustment are 0.100 and 0.122. The apparent peak at $q = 0.04$ is driven by a subset of replications with many small p-values.

Note that here, the inflated proportion of false discoveries is a consequence of correlation through shared factors, varying infection intensity, among other things, and not a consequence of pre-labeling strategy.

| Pre-labels | $q$ | Balanced accuracy | DE F1 | Null $p \leq 0.05$ |
|---|---|---|---|---|
| Random | 0.01 | 0.720 (0.026) | 0.039 (0.009) | 0.1076 (0.0186) |
| Viral load | 0.01 | 0.866 (0.010) | 0.039 (0.012) | 0.0625 (0.0075) |
| Random | 0.04 | 0.864 (0.016) | 0.054 (0.014) | 0.1652 (0.0296) |
| Viral load | 0.04 | 0.843 (0.016) | 0.054 (0.010) | 0.1857 (0.0317) |
| Random | 0.10 | 0.880 (0.011) | 0.058 (0.012) | 0.1472 (0.0248) |
| Viral load | 0.10 | 0.838 (0.010) | 0.055 (0.011) | 0.1372 (0.0256) |

**Table R1.** Model 1 results across pre-labeling proportions at $f = 2$. Infection balanced accuracy is evaluated among unlabeled test cells. DE F1 concerns identification of infection-associated genes. The rightmost column is the fraction of raw p-values at or below 0.05 among tested null gene–cell-type pairs outside the training panel. Each entry is mean (standard error) over 50 replications.

##### 2.2.2 Ambiguous boundary cells

Under Model 1, the infection effect in cell $i$ is multiplied by its intensity $\xi_i$. Thus, an infected cell with small $\xi_i$ is, from the perspective of our model, more like an uninfected cell. We divided the realized $\xi_i \in (0, 1)$ into the fixed intervals $[0, .25]$, $(.25, .50]$, $(.50, .75]$, and $(.75, 1]$. In Figure R2, we display sensitivity across these intervals at $f \in \{1.5, 2, 4\}$ and $q = 0.04$, with viral-load pre-labeling. To clarify, sensitivity is the fraction of truly infected, unlabeled test cells in an interval that are classified as infected.

We can see that with viral-load pre-labeling, sensitivity rises from 0.163 in the lowest intensity interval to 0.948 in the highest at $f = 2$. The same relative performances appear at the other fold changes.

##### 2.2.3 Combinations of model assumption violations under negative binomial

Next, we varied the log-library-size standard deviation $\sigma_U$, the nonzero factor-loading standard deviation $\sigma_L$, and the infection-intensity parameter $a$. When intensity does not vary, we set $B_i = 1$ and denote this setting by $a = \infty$. The parameter $\pi_V$ is the fraction of prelabels chosen by highest viral count among all cells: we only consider 0 and 1 (i.e., none or all). Table R2 lists the displayed $(\sigma_U, \sigma_L, a, \pi_V)$ combinations. We simulated by varying the

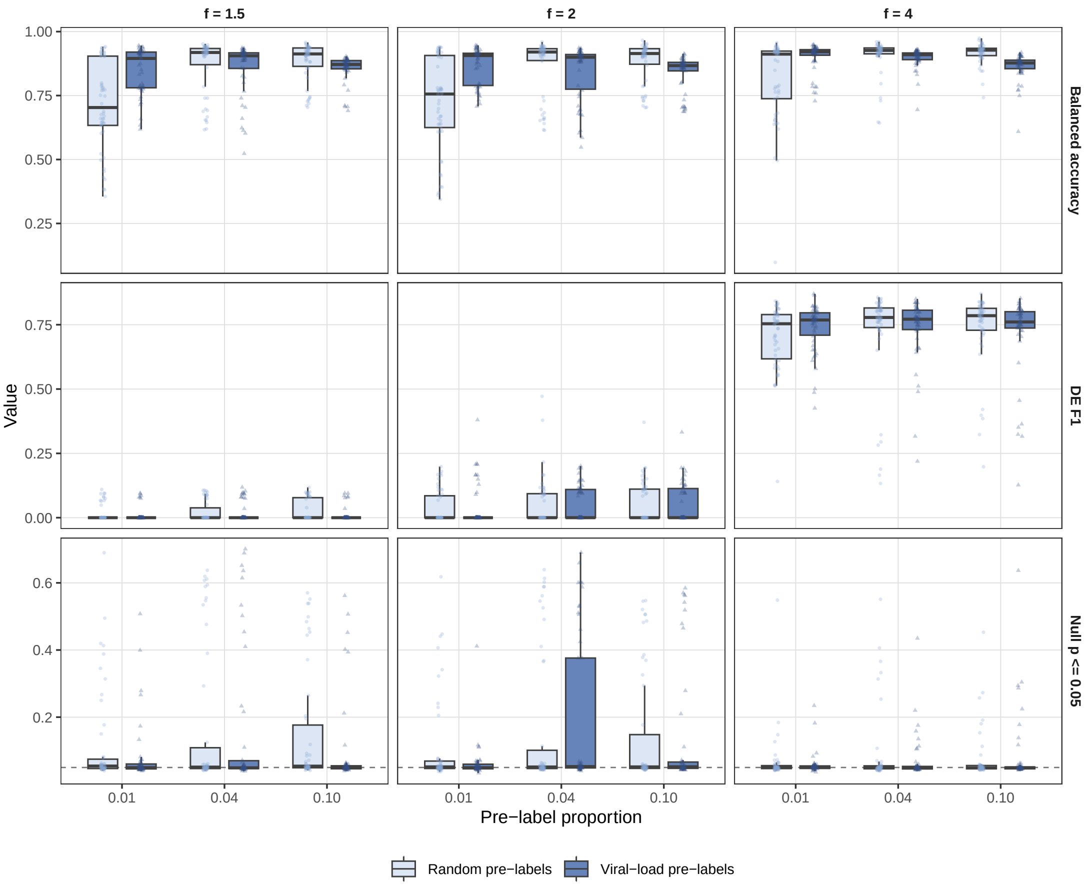

**Fig. R1.** Model 1 results across pre-labeling proportions. Infection balanced accuracy (top row), DE F1 (middle row), and the fraction of raw p-values at or below 0.05 among tested null gene–cell-type pairs outside the training panel (bottom row), against pre-label proportion. Columns indicate infection fold change. Boxes show the median and interquartile range across 50 replications; whiskers extend to 1.5 times the interquartile range, and faint points show individual replications. The dashed line in the bottom row marks the nominal 0.05 fraction.

parameters $\sigma_U \in \{0, 0.35, 0.55\}$, $\sigma_L \in \{0, 0.35, 0.55\}$, or $a \in \{\infty, 2\}$. All scenarios were run using scDEcrypter's default settings.

The proportion of truly null genes not included in the training set having a p-value $< 0.05$ was (on average) 0.0508 at baseline (assuming independent counts), 0.0514 with library-size variation, and 0.0516 when factor-loading standard deviation 0.35 was added (i.e., under mild correlation). Introducing varying intensity with the same library-size variation but no factors raised it to 0.108. With the additional factors inducing correlation, it

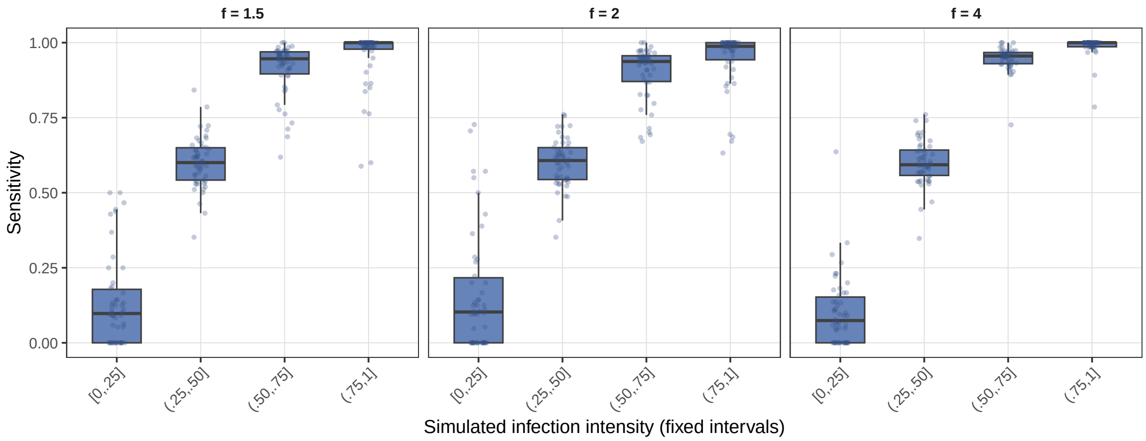

**Fig. R2.** Sensitivity under Model 1 across infection-intensity intervals for three infection fold changes at $q = 0.04$ with viral-load pre-labeling.

reached 0.165.

Repeating the six settings with viral-load labels changed little when the infection was simulated as binary. With variable intensity, the non-training proportion increased from 0.108 to 0.131 without factors and from 0.165 to 0.186 with factors. Thus, variable intensity worsens calibration in this design, and factors amplify it.

| Count setting | $(\sigma_U, \sigma_L, a, \pi_V)$ | Other null | Training null | BH FDP | Balanced accuracy |
|---|---|---|---|---|---|
| Independent counts | $(0, 0, \infty, 0)$ | 0.0508 (0.0004) | 0.0517 (0.0030) | 0.0442 (0.0026) | 1.0000 (0.0000) |
| Library size only | $(.35, 0, \infty, 0)$ | 0.0514 (0.0007) | 0.0539 (0.0036) | 0.0485 (0.0028) | 0.9999 (0.0001) |
| Sparse factors | $(.35, .35, \infty, 0)$ | 0.0516 (0.0009) | 0.0528 (0.0030) | 0.0465 (0.0029) | 1.0000 (0.0000) |
| Stronger factors | $(.35, .55, \infty, 0)$ | 0.0556 (0.0023) | 0.0907 (0.0139) | 0.1024 (0.0241) | 0.9867 (0.0050) |
| Variable intensity | $(.35, 0, 2, 0)$ | 0.1076 (0.0227) | 0.1500 (0.0372) | 0.1476 (0.0459) | 0.9013 (0.0089) |
| Intensity and factors | $(.35, .35, 2, 0)$ | 0.1652 (0.0296) | 0.2607 (0.0508) | 0.2462 (0.0580) | 0.8644 (0.0160) |
| Independent counts | $(0, 0, \infty, 1)$ | 0.0513 (0.0004) | 0.0482 (0.0026) | 0.0426 (0.0019) | 1.0000 (0.0000) |
| Library size only | $(.35, 0, \infty, 1)$ | 0.0506 (0.0008) | 0.0536 (0.0036) | 0.0453 (0.0026) | 1.0000 (0.0000) |
| Sparse factors | $(.35, .35, \infty, 1)$ | 0.0508 (0.0009) | 0.0513 (0.0034) | 0.0432 (0.0038) | 1.0000 (0.0000) |
| Stronger factors | $(.35, .55, \infty, 1)$ | 0.0564 (0.0022) | 0.0902 (0.0137) | 0.0950 (0.0202) | 0.9889 (0.0047) |
| Variable intensity | $(.35, 0, 2, 1)$ | 0.1309 (0.0268) | 0.1874 (0.0438) | 0.1795 (0.0512) | 0.8759 (0.0113) |
| Intensity and factors | $(.35, .35, 2, 1)$ | 0.1857 (0.0317) | 0.2892 (0.0523) | 0.2809 (0.0591) | 0.8426 (0.0155) |

**Table R2.** Model 1 results across selected settings varying $(\sigma_U, \sigma_L, a, \pi_V)$. $a = \infty$ means constant infection intensity among infected cells. The upper and lower blocks repeat the same six count settings with $\pi_V = 0$ and $\pi_V = 1$. Null columns give the proportion of known-null p-values at or below 0.05, partitioned over genes included in the generation set (training) and those note (other). BH FDP is the false discovery proportion after BH adjustment. Entries represent the mean (standard error) over 50 replications.

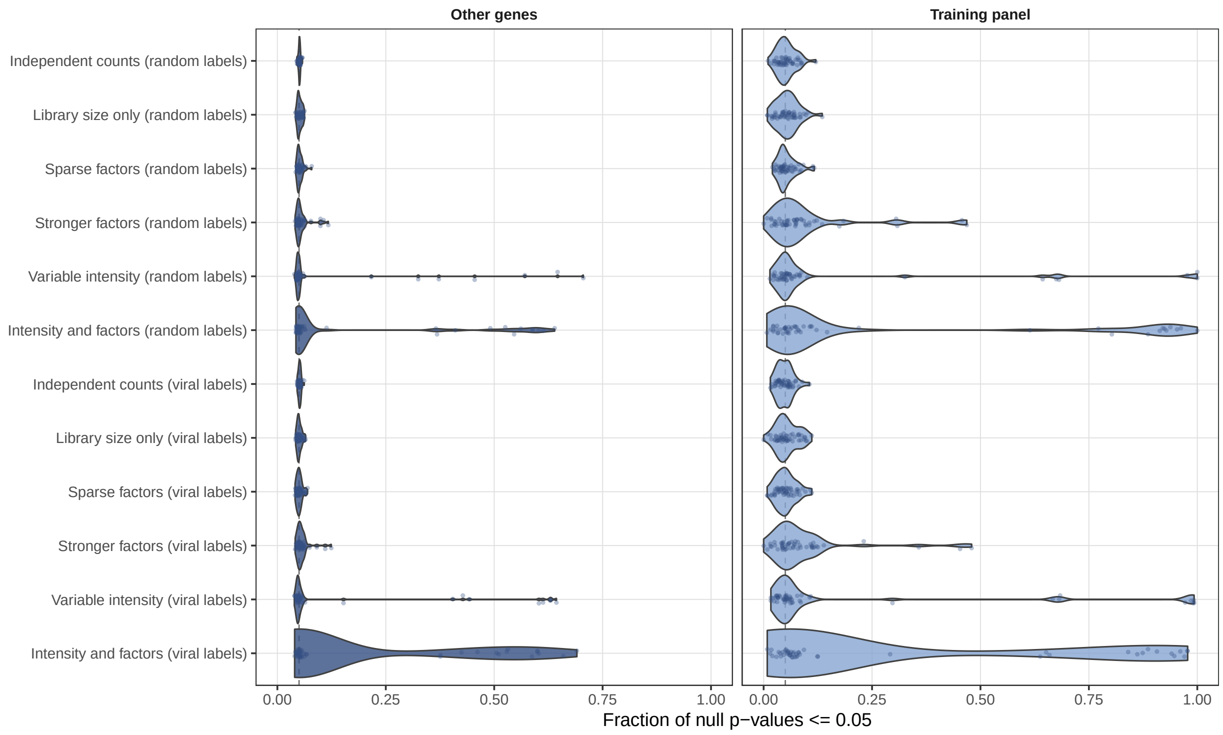

**Fig. R3.** Known-null p-values at or below 0.05 across count-model departures for Model 1. The left panel indicates known-null genes not included in the training set and the right panel are known-null genes used for training. Each violin shows the distribution across 50 replications. The dashed line marks 0.05.

##### 2.2.4 Genes used to estimate the weights

In Table R3, we split genes into panel versus non-panel (other) under Model 1 with $f = 2$ and $q = 0.04$. Training panel genes are those used to train the model and estimate the weights. Figure R4 shows the fraction of known-null p-values at or below 0.05 in each replication under random or viral-load pre-labeling with $q = 0.04$ and $f \in \{1.5, 2, 4\}$.

In the most difficult case considered, Model 1 with $(\sigma_U, \sigma_L, a, \pi_V) = (.35, .35, 2, 1)$, the fraction of known-null raw p-values at or below 0.05 was 0.186 for other genes and 0.289 for panel genes on average. These numbers are slightly skewed by a small number of replicates with a high proportion of small p-values. Notably, both groups were near 0.05, on average, in the independent count baseline. Recomputing the weights during training-gene permutations preserved gene–weight dependence, but cannot repair departures from the exchangeability assumption.

| $(\sigma_U, \sigma_L, a, \pi_V)$ | Genes | $p \leq 0.01$ | $p \leq 0.05$ |
|---|---|---|---|
| $(0, 0, \infty, 0)$ | Other | 0.0104 (0.0002) | 0.0508 (0.0004) |
| $(0, 0, \infty, 0)$ | Training | 0.0115 (0.0013) | 0.0517 (0.0030) |
| $(.35, .35, 2, 1)$ | Other | 0.1009 (0.0217) | 0.1857 (0.0317) |
| $(.35, .35, 2, 1)$ | Training | 0.2337 (0.0504) | 0.2892 (0.0523) |

**Table R3.** False discovery proportions under independent counts vs dependent counts with varying infection intensity under Model 1 with $f = 2$. Entries are mean (standard error) proportions of known-null p-values below each threshold over 50 replications.

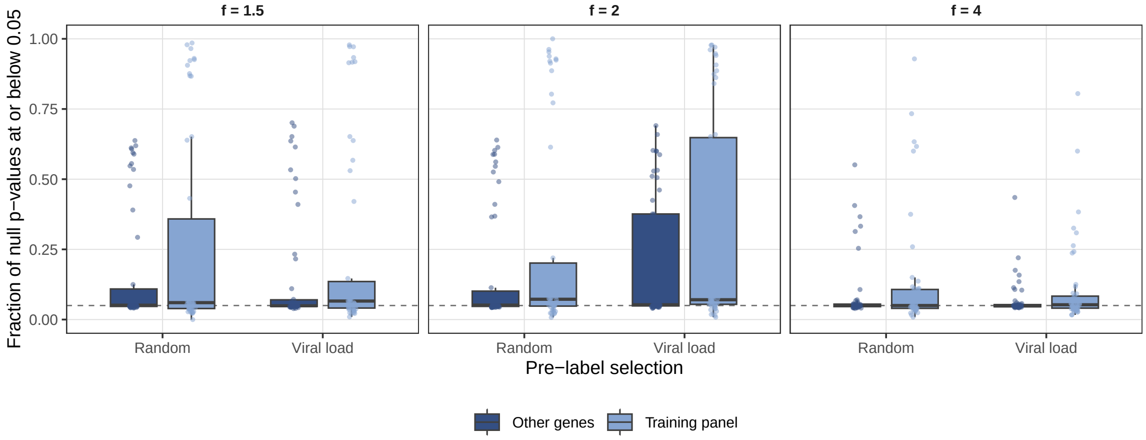

**Fig. R4.** Proportion of known-null p-values at or below 0.05 under Model 1 with $q = 0.04$ and $(\sigma_U, \sigma_L, a, \pi_V) = (.35, .35, 2, 1)$. Each boxplot shows 50 replication-level fractions, averaged across cell types within each replication. The dashed line marks 0.05. The $f = 2$, viral-load boxes correspond to the bottom rows of Table R3. The independent-count baseline appears only in the table. The $f = 2$ viral-load means are reported in Table R3.

##### 2.2.5 Explicit gene correlation

As described, under Model 2, we varied the AR(1) correlation $\rho$ and infection fold change $f \in \{1.5, 2, 4\}$, with $q = 0.04$. Table R4 reports known-null genes outside the training panel. Figure R5 also shows training-panel known-null genes. The training panel was selected as the 34 most variable genes per cell type on the generation set cells. At $f = 4$, only infection-associated genes ended up in the training panel for all 50 replications at each correlation, so panel-gene null calibration was undefined.

The proportion of known-null p-values at or below 0.05 stayed near 0.05 through $\rho = 0.95$. At $\rho = 0.99$ it reached 0.085 for $f = 1.5$, while remaining near 0.05 for $f = 2$ and $f = 4$. At $\rho = 0.999$ it reached 0.327 and 0.224 for $f = 1.5$ and $f = 2$, respectively, but remained 0.052 for $f = 4$. Thus, very strong local correlation disrupts calibration when infection signal is weaker.

| AR(1) correlation | $f = 1.5$ | $f = 2$ | $f = 4$ |
|---|---|---|---|
| $\rho = 0.3$ | 0.0503 (0.0003) | 0.0500 (0.0003) | 0.0497 (0.0004) |
| $\rho = 0.5$ | 0.0498 (0.0003) | 0.0492 (0.0003) | 0.0491 (0.0003) |
| $\rho = 0.7$ | 0.0513 (0.0003) | 0.0494 (0.0003) | 0.0491 (0.0004) |
| $\rho = 0.9$ | 0.0531 (0.0005) | 0.0494 (0.0004) | 0.0496 (0.0005) |
| $\rho = 0.95$ | 0.0560 (0.0009) | 0.0493 (0.0007) | 0.0495 (0.0007) |
| $\rho = 0.99$ | 0.0850 (0.0023) | 0.0517 (0.0016) | 0.0514 (0.0015) |
| $\rho = 0.999$ | 0.3271 (0.0129) | 0.2241 (0.0285) | 0.0516 (0.0046) |

**Table R4.** Model 2 across $\rho$ settings. Proportion of known-null p-values at or below 0.05 among genes outside the training panel under AR(1) Gaussian correlation. Entries are mean (standard error) over 50 replications.

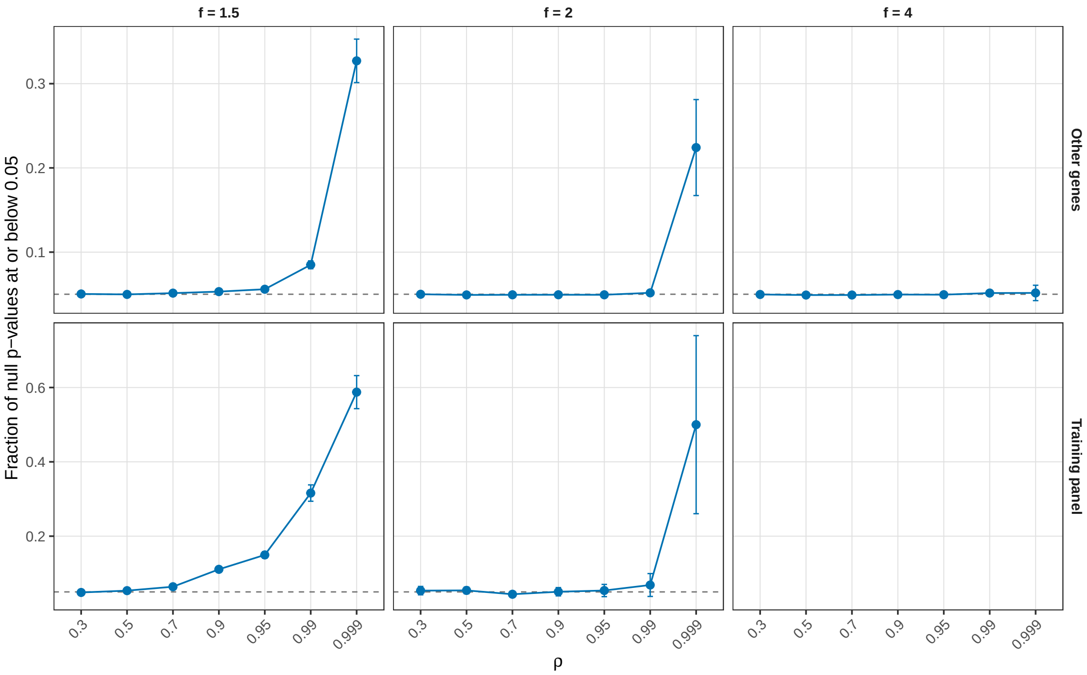

**Fig. R5.** Model 2 across $\rho$ and $f$ settings. Proportion of known-null p-values at or below 0.05 under AR(1) Gaussian correlation, for other genes (top) and training-panel genes (bottom). Points are means across replications with available null genes; bars span two standard errors. All plotted groups have 50 replications except training-panel genes at $f = 2$, where $\rho = 0.99$ has 47 and $\rho = 0.999$ has 14. No training-panel null genes were available at $f = 4$. The dashed line marks 0.05.

#### 3 Supplementary Figures

Figures S1 and S2 summarize additional simulation results, and Figure S3 reports computational benchmarks. Figures S4 and S5 provide additional influenza results; Figures S6 and S7 provide additional SARS-CoV-2 results.

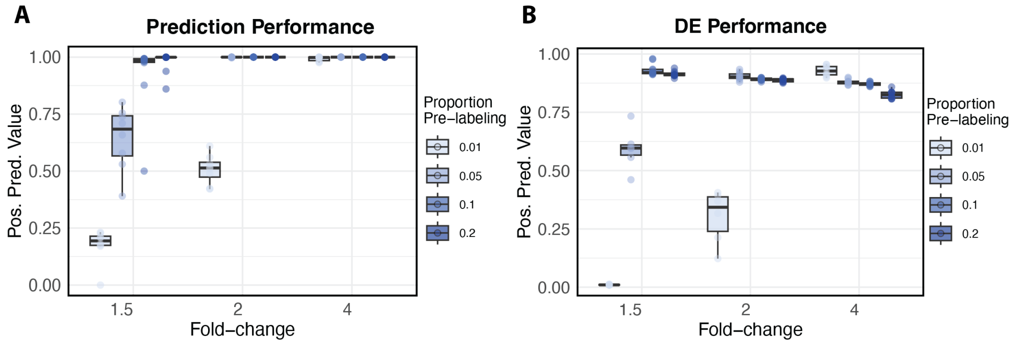

**Fig. S1.** Positive predictive value across simulation settings, evaluated separately for infection-state prediction and DE testing. A. Positive predictive value for infection-state thresholding as a function of effect size (fold-change) and proportion of cells pre-labeled for model fitting. B. Positive predictive value for differential expression testing under the same simulation settings. In both panels, boxplots summarize the distribution of positive predictive value across simulation replicates, with individual points overlaid and colored by the proportion of pre-labeled cells.

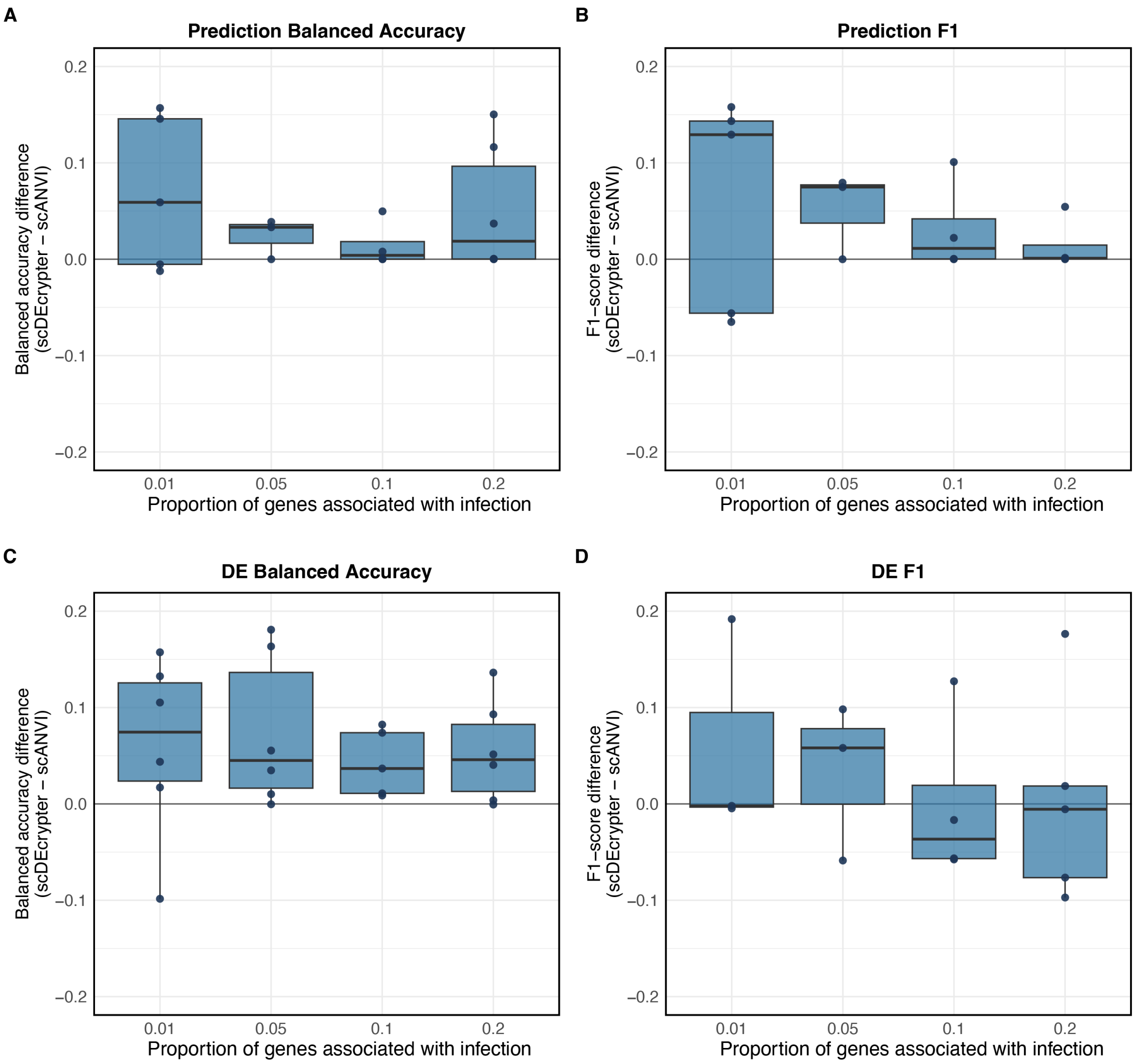

**Fig. S2.** Comparison of scDEcrypter and scANVI across simulated scenarios. Difference in performance between scDEcrypter and scANVI was calculated as scDEcrypter minus scANVI; values above zero favor scDEcrypter. A, B. Balanced accuracy (A) and F1 (B) for infection-state prediction. C, D. Balanced accuracy (C) and F1 (D) for DE testing. Points denote simulation scenarios averaged over replicates; the horizontal line marks equal performance.

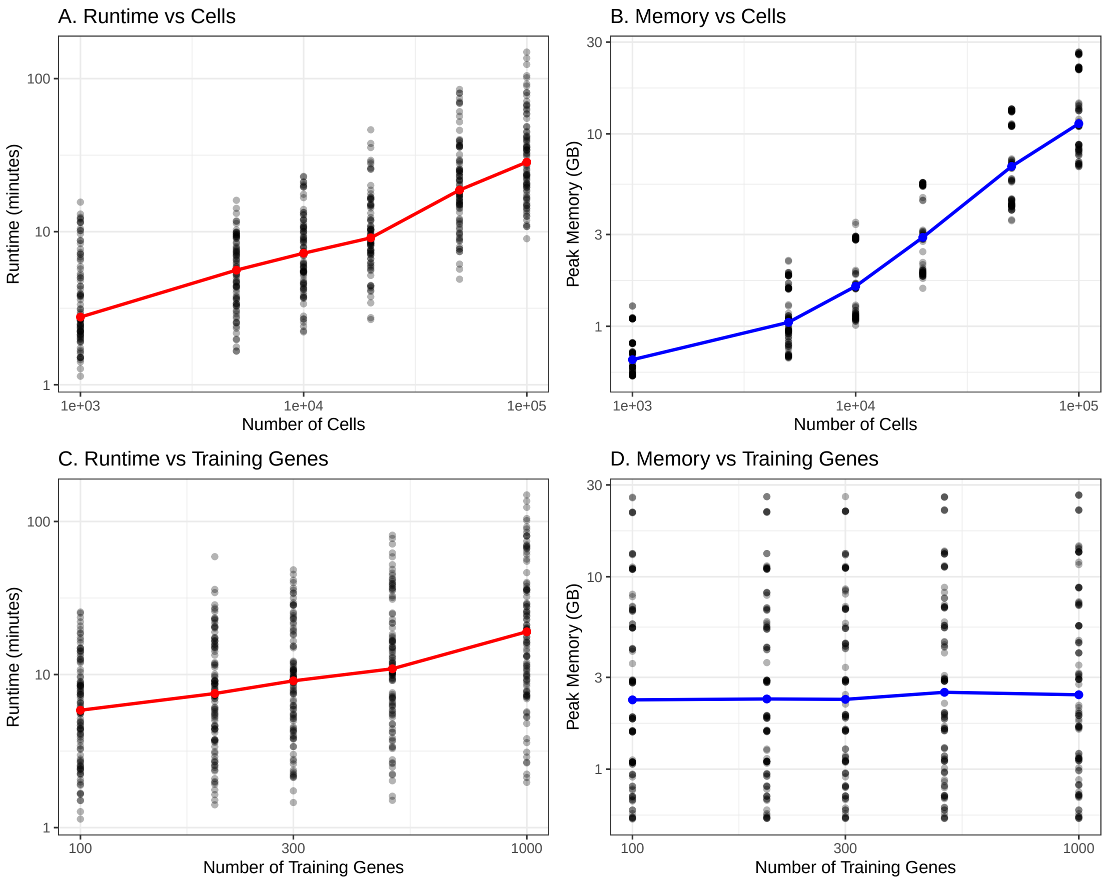

**Fig. S3.** Computational benchmarking of scDEcrypter. A. The total runtime (minutes) of scDEcrypter training and testing on datasets ranging from 1,000 to 100,000 cells. B. The peak memory usage (GB) during scDEcrypter training and testing across dataset sizes. C, D. Runtime (C) and peak memory usage (D) across the number of genes used for training.

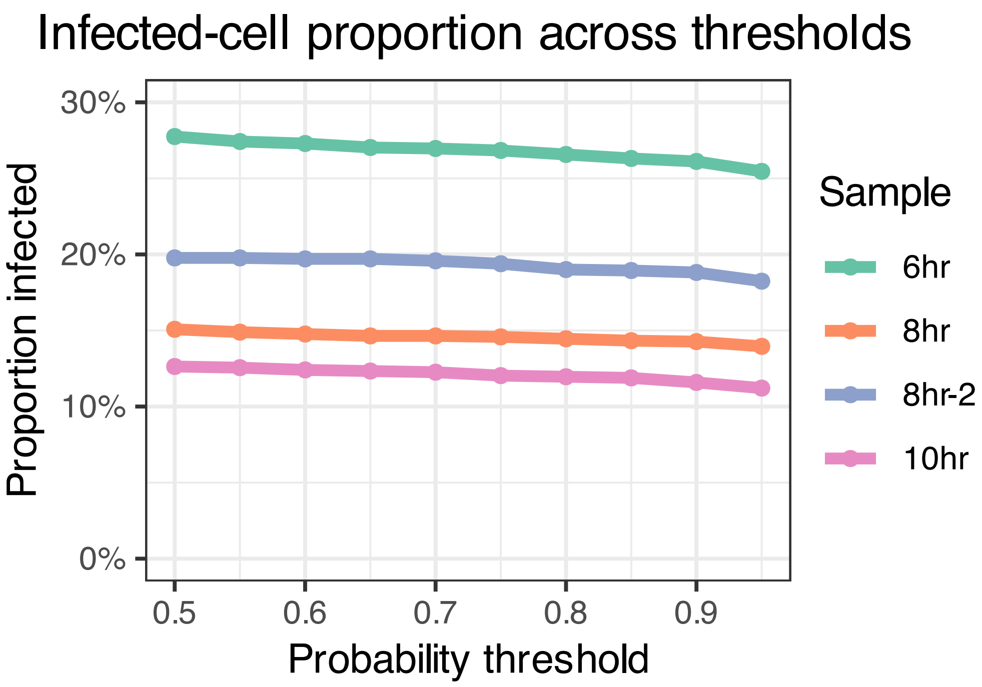

**Fig. S4.** Infection state prediction on the influenza dataset as a function of the probability weight threshold.

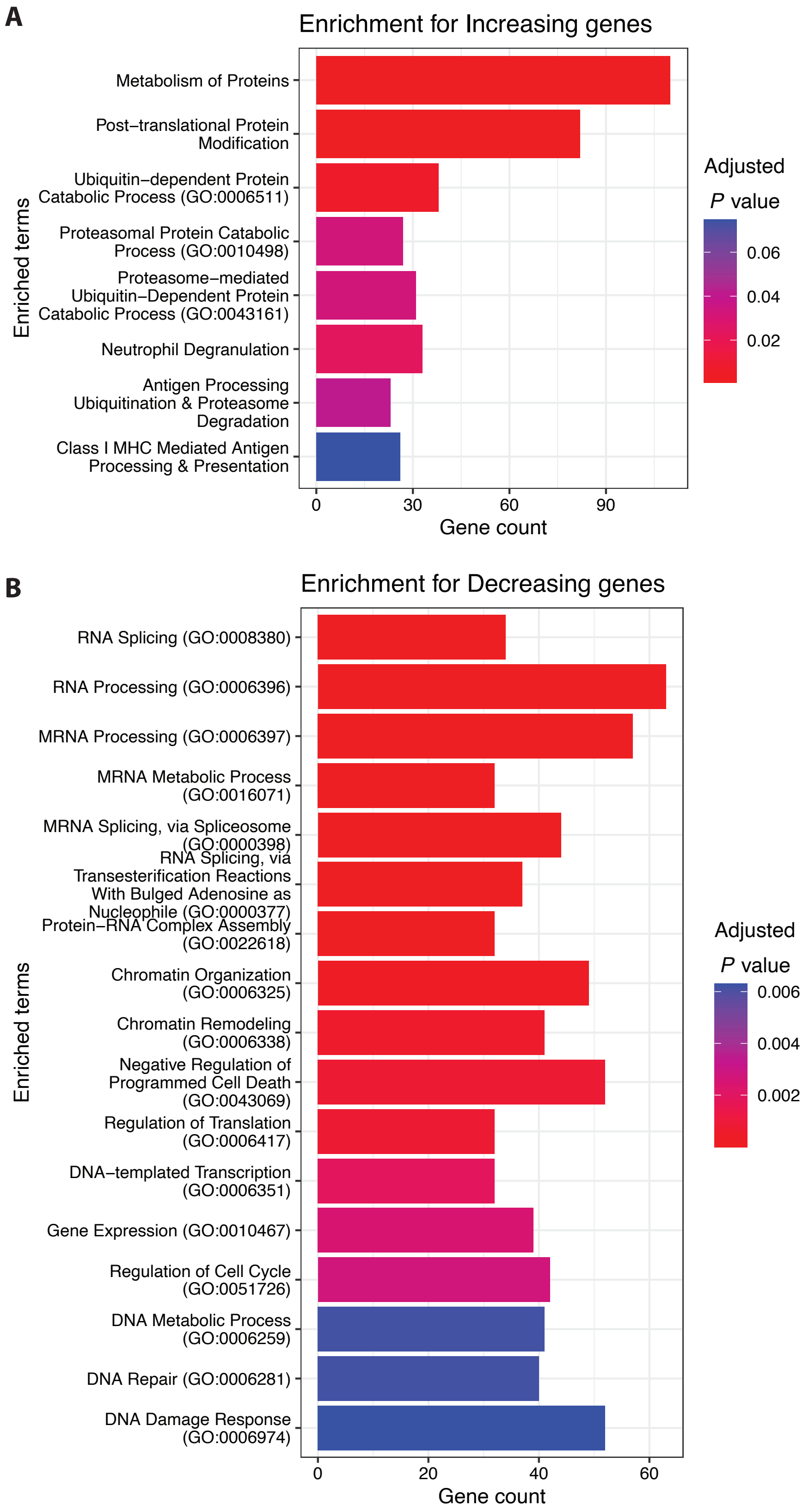

**Fig. S5.** GO Biological Processes (2026) enrichment among DE genes in the influenza dataset with progressively increasing (A) or decreasing (B) expression in the infected state over time.

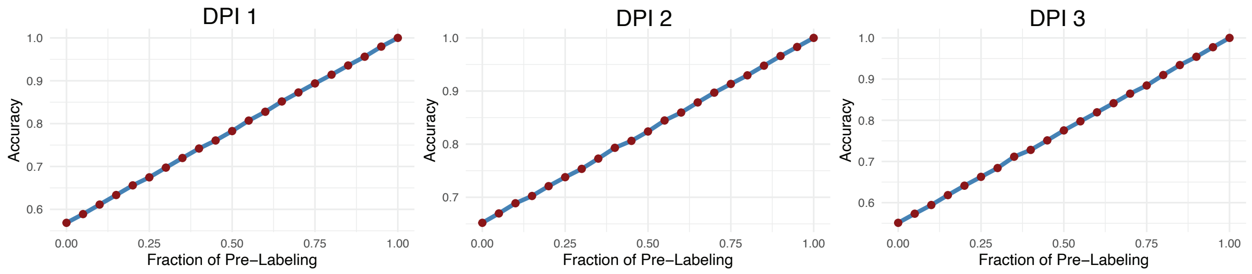

**Fig. S6.** Overall accuracy in cell-type prediction in the SARS-CoV-2 dataset as a function of pre-labeling in the test dataset in the one day post-infection (DPI) sample (left), two days (middle), and three days (right).

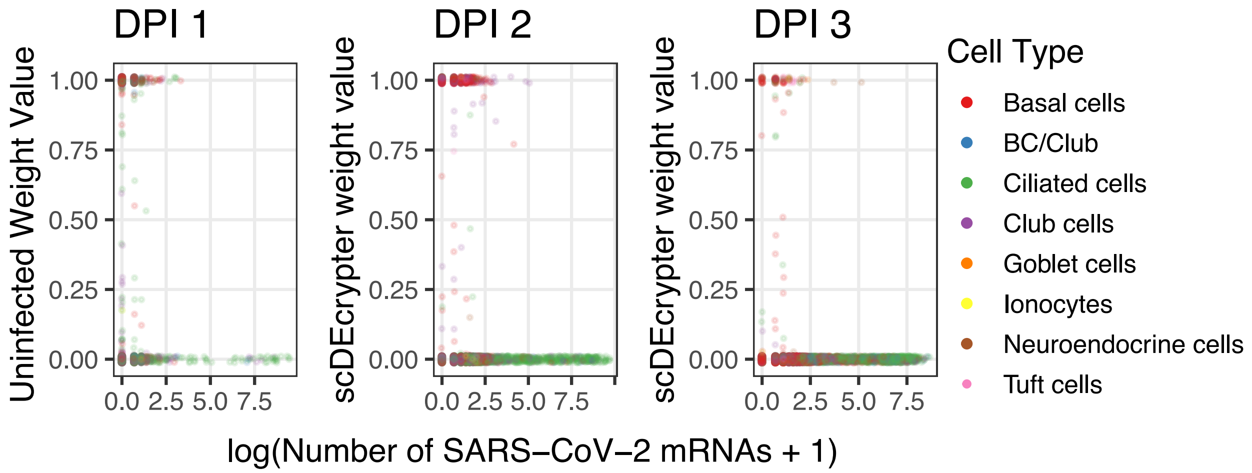

**Fig. S7.** scDEcrypter-derived probability of the uninfected cell state as a function of viral load. The pre-existing infection label was cleared for these cells, and the fitted mixture model (trained on an independent generation set) was applied to the test set to estimate each cell's posterior probability of belonging to the uninfected state. The x-axis shows the log-transformed total number of SARS-CoV-2 mRNAs detected per cell; the y-axis shows the model-estimated probability of the uninfected state. Points are colored by annotated cell type.

### References

Jinwen Fu, Aaron J Molstad, and Hui Zou. A direct approach to tree-guided feature aggregation for high-dimensional regression. *arXiv preprint arXiv:2507.19650*, 2025.

Aaron J Molstad and Keshav Motwani. Multiresolution categorical regression for interpretable cell-type annotation. *Biometrics*, 79(4):3485–3496, 2023. Publisher: Wiley Online Library.
